# Implicit micelle model for membrane proteins using super-ellipsoid approximation

**DOI:** 10.1101/654103

**Authors:** Takaharu Mori, Yuji Sugita

## Abstract

Surfactant micelles are often utilized as membrane mimetics for structure determination and functional analysis of membrane proteins. Although curved-surface effects of the micelle can perturb their structure, it is difficult to assess such effects and membrane mimetic artifacts by experimental and theoretical methods. Here, we propose an implicit micelle model (IMIC) to be used in molecular dynamics (MD) simulations of membrane proteins. IMIC is an extension of the IMM1 implicit membrane model by introducing a super-ellipsoid approximation to represent the curved-surface effects. Most of the parameters for IMIC are obtained from all-atom explicit solvent MD simulations of twelve membrane proteins in various micelles. In simulations of the HIV envelop protein gp41, M13 major coat protein gp8, and amyloid precursor protein (APP) dimer, curved-surface and compact hydrophobic-core effects are exhibited. The MD simulations with IMIC provide accurate structure predictions of membrane proteins in various micelle environments quickly with smaller computational cost than that necessary for explicit solvent/micelle model.

## INTRODUCTION

Membrane proteins play important roles in cellular processes like substrate transport, signal transduction, membrane fusion, and cell adhesion. Cell membranes are mainly composed of phospholipids, which spontaneously assemble into bilayers, with hydrophilic head groups exposed to water and hydrophobic tail groups buried inside the membrane. Experiments on membrane proteins frequently use surfactant as a membrane mimetic agent to purify and stabilize the proteins, where the surfactant covers the transmembrane (TM) domain by forming a micelle.^1^ X-ray crystallography often detects electron densities of bound surfactants,^2^ and recent cryo-electron microscopy (cryo-EM) experiments clearly discern an ellipsoidal micelle around the TM domain.^3-6^ In solution NMR, proteins are frequently reconstituted into micelles, bicelles, or nanodiscs,^7^ while in solid-state NMR they are in multilamellar vesicles or oriented lipid bilayers.^8,9^

It is commonly recognized that the surfactant micelle may not mimic a lipid bilayer environment faithfully, in that, compared to lipid bilayers, the micelle has a highly curved surface and compact hydrophobic core region.^10,11^ Many membrane proteins have been shown to have different conformations in micelles and bilayers. Examples include the LLP-3 domain in the HIV-1 envelope glycoprotein gp41,^12^ bacteriophage M13 major coat protein gp8,^13^ influenza M2 proton channel,^14^ phospholamban,^15,16^ epidermal growth factor receptor (EGFR),^17^ and amyloid precursor psorein (APP).^18-20^ Specifically, the LLP-3 peptide forms a curved α-helix on the micelle surface, but is straight on the bicelle surface.^12^ The M13 major coat protein gp8 forms various kinked-helix conformations in micelles, while a continuous α-helix predominates in fully hydrated vesicles.^13^ This is in contrast to the glycophorin A dimer, the conformation of which is identical in micelles and lipid bilayers, presumably because the two monomers are tightly bound through GxxxG motif interactions.^21^ Therefore, among experimentally determined structures, discrimination between native structures and membrane mimetic artifacts is extremely valuable.

Molecular dynamics (MD) simulation has been useful for investigation of the structural and dynamic properties of membrane proteins in surfactant micelles as well as in lipid bilayers.^22-25^ Recent development of automated system builders, such as CHARMM-GUI *Micelle builder*,^26^ makes it easy to model a protein-micelle complex for all-atom MD simulations in explicit solvent and micelle. One difficulty, though, is that the aggregation number of surfactant around the protein is usually unknown, and there is limited experimental data available in this regard.^27-30^ To avoid this problem, protocols for the spontaneous formation of the protein-micelle complex have been proposed, where the MD simulation starts from a random distribution of surfactants in the simulation box.^31-33^ Since such simulations require long relaxation times (typically, several hundred nanoseconds), course-grained models or implicit water models have been employed to enhance surfactant self-assembly.^34-37^ Even if the optimal aggregation number of surfactant could be determined experimentally or theoretically, there is large computational cost for simulation of a protein-micelle complex in explicit solvent.

The implicit solvent model can significantly reduce computational cost in biomolecular simulations.^38^ In this model, the solvation free energy of solute is incorporated into a molecular mechanics potential energy function as the effective energy term. The solvation free energy is traditionally decomposed into electrostatic and nonpolar contributions. The electrostatic term is computed by numerically solving the Poisson-Boltzmann (PB) equation,^39^ or by using screened pairwise interactions between partial charges of solute atoms based on the Generalized Born theory (GB).^40^ The nonpolar term can be calculated from the solvent accessible surface area (SASA).^41^ In other methods such as hydration shell models, the solvation free energy is assumed to be proportional to the solvent accessible volume of the first hydration shell.^42^ In the EEF1 model proposed by Lazaridis and Karplus,^43^ the solvent excluded volume is estimated from the volume of neighboring solute atoms.

Implicit solvent models have been also extended to the membrane environment,^44^ and most of them are based on the GB method.^45-49^ They introduce a low dielectric slab (*ε* = 1-4) at *z* = 0, and parameters are optimized to reproduce PB calculations or all-atom MD simulations. The EEF1 model was also extended to the membrane model (IMM1), where the solvation free energy is calculated by a combination of the solvation free energies in water and cyclohexane.^50^ The IMM1 model has been further extended to anionic lipid bilayers,^51^ mixed-lipid bilayers,^52^ and bilayers with a membrane potential,^53^ aqueous pore,^54^ and large curvature.^55^ Although many implicit lipid bilayer models have been developed, to our knowledge “micelle models” have not been proposed.

In this study, we propose a new implicit solvent model, named the IMIC model, which treats the micelle environment implicitly. It is an extended form of the IMM1 model. We introduce a super-ellipsoid approximation to define micelle shape, and the parameters are derived from all-atom MD simulations of pure micelles and protein-micelle complexes in explicit solvent. We validate the IMIC model by comparing structures and dynamics of selected membrane proteins with those in all-atom MD simulations. In application studies, we simulate the HIV-1 envelope glycoprotein gp41, bacteriophage M13 major coat protein gp8, and APP dimer to examine whether the IMIC/IMM1 models identify membrane mimetic artifacts in solution NMR structures in micelles.

## THEORY

### Approximation of micelle shape

First, we propose a “super-ellipsoid” approximation to define micelle shape. For simplicity, the micelle is centered at the origin of the system (*x, y, z*) = (0, 0, 0). A pure micelle has been typically approximated as an “ordinary” ellipsoid.^56-58^ As observed in recent cryo-EM experiments,^3-6^ micelle shape can be deformed in the presence of membrane proteins, resulting in a flat region near the micelle center where the protein is contained. Both simple and deformed shapes can be described with the super-ellipsoid function:

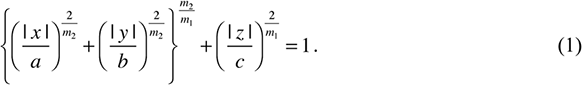

*a, b*, and *c* are the semi-axes of the super-ellipsoid, and *m*_1_ and *m*_2_ determine the shape of the cross section in the super-ellipsoid (see Figure S1).^59,60^ In the case of *m*_1_ = *m*_2_ = 1, eq 1 gives an ordinary ellipsoid. If 0 < *m*_1_ < 1 and *m*_2_ = 1, the cross section in a plane perpendicular to the *XY*-plane is expanded, keeping the semi-axes at the given lengths, and the shape also resembles a bicelle or nanodisc. If *m*_1_ = 1 and 0 < *m*_2_ < 1, the cross section in a plane parallel to the *XY*-plane is expanded. The shape becomes close to rectangle as both *m*_1_ and *m*_2_ decrease. Note that *m* < 0 or *m* > 1 is not allowed, because it produces a non-micelle-like shape resembling an octahedron.

### Implicit micelle model (IMIC)

The implicit micelle model (IMIC) creates an ellipsoidal hydrophobic core region in the system using the super-ellipsoid function. The IMIC model is an extension of the EEF1/IMM1 models,^43,50^ where the effective energy *W* of a solute molecule is defined as the sum of molecular mechanics potential energy *E*_MM_ and solvation free energy Δ*G*^solv^, given by

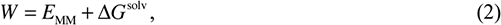

Where

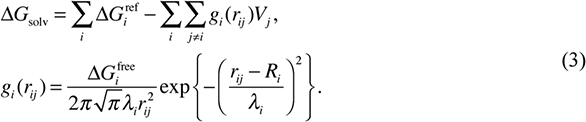

*r*_*ij*_ is the distance between atoms *i* and *j*, and *V*_*j*_ is the volume of the *j*-th atom. The function *g*_*i*_ is the density of the solvation free energy of the *i*-th atom, defined with the van der Waals radius *R*_*i*_ and thickness of the first hydration shell *λ*_*i*_ · 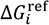 is the solvation free energy of the atom when it is fully exposed to solvent. 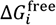 is similar to 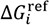, but is determined to satisfy the zero solvation energy of deeply buried atoms.^43^

As in the IMM1 model, 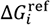 is defined as a combination of the solvation free energies of the *i*-th solute atom in water and cyclohexane:

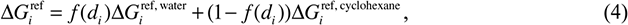

where *f*(*d*) is a function that describes the transition between water and cyclohexane phases. In the IMIC model, we use the following function:

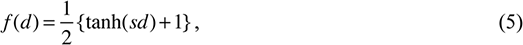

where *d* is the depth of the solute atom from the super-ellipsoid surface (Figure 1a), and *s* controls the steepness of the micelle-water interface. If a solute atom is inside the micelle, *d* has a negative value, and vice versa. *d* is computed with an iterative minimization scheme (see Supporting Information).^61^ The function *f* produces a sigmoidal curve, and it gives 0.5 at the super-ellipsoid surface (*d* = 0) (Figure 1b). In our model, we assume that the surfactant headgroup has the same properties as aqueous solution,^50^ and thus, the midpoint *f* = 0.5 corresponds to the boundary between surfactant hydrocarbon and polar headgroups. The derivatives of the solvation free energy with respect to atomic position are also calculated (see Supporting Information).

**Figure 1.**
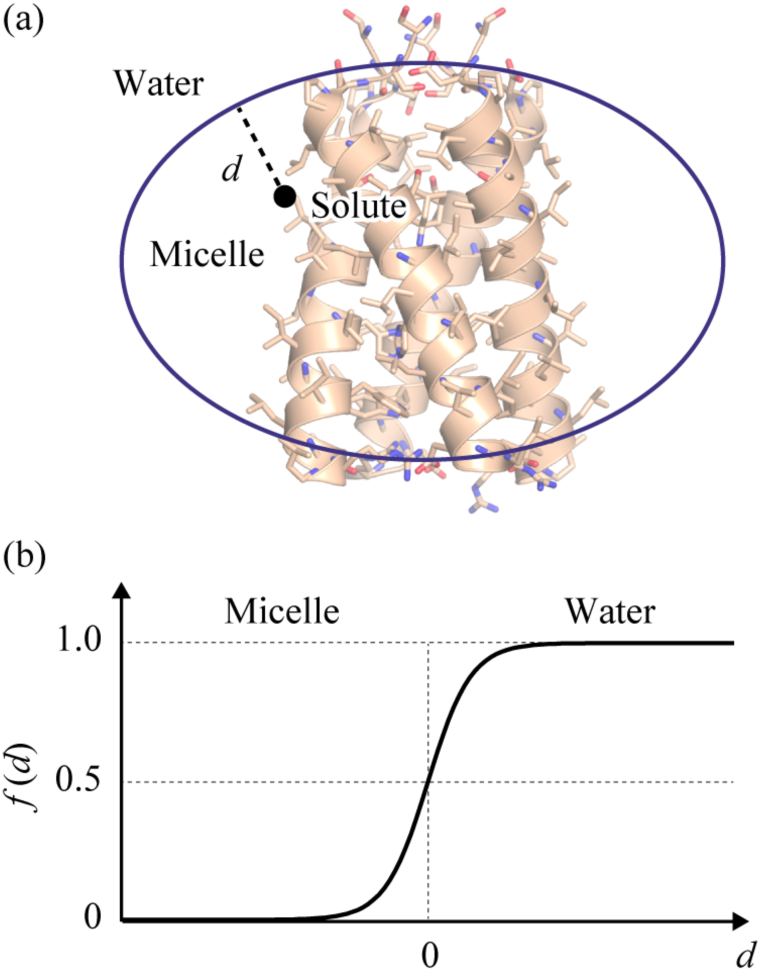
Schematic representation of a membrane protein in the IMIC model. (a) The depth *d* of a solute atom is defined as the minimum distance from the super-ellipsoid surface. (b) Sigmoidal curve of the function *f*(*d*) in eq 5.

As in the IMM1 model, we use a distance-dependent dielectric constant for electrostatic interactions. The dielectric constant depends on the positions of interacting atoms,^50^ defined as

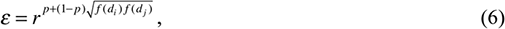

where *r*_*ij*_ is the distance between the *i*-th and *j*-th atoms, and *p* is an empirical parameter to adjust strength of the interactions (*p* = 0.85 for CHARMM19^50^ and 0.91 for CHARMM36^62,63^) Far from the micelle surface, the dielectric constant *ε* is close to *r*, corresponding to the EEF1 model, while in the micelle center, it provides strengthened interactions. Eqs 1 and 5 are the only differences between the IMIC and IMM1 models. The IMM1 model utilizes *f* (*z*′) = *z*′^*n*^/(1 + *z*′^*n*^) for eq 5, where *z*′ = |*z*|/(*T/2*) and *T* is the membrane thickness. The IMIC model is nearly equivalent to the IMM1 model when *a* and *b* → ∞ and *c* is half membrane thickness.

We implemented the IMIC, EEF1, and IMM1 models into GENESIS,^64,65^ and these were parallelized with hybrid MPI/OpenMP protocols. Computational cost of the IMIC model is ∼1.9-times that of the vacuum simulations, and ∼1.2-times that of IMM1 simulations, which is mainly due to the iterative minimization scheme for the computation of *d*. In this study, we use the IMM1-p36 parameter set for Δ*G*ref, Δ*G*free, *λ*, and *V*, which is an updated version of the CHARMM19-based IMM1 model for the CHARMM C36 force fields.^62^ We used top_all36_prot_lipid_eef1.1.rtf and solvpar22.inp for topology and parameter files, respectively, both of which are available in the CHARMM program package.^66^

## METHODS

In order to carry out MD simulations of membrane proteins in the IMIC model with a given number of surfactant molecules, we should estimate the micelle size and shape in advance. Let us consider a structural change of the micelle upon protein insertion (Figure 2), where the surfactant aggregation number is assumed to be invariant. If the protein is inserted into the micelle along the *Z*-axis as in lipid bilayers, the semi-axis *c* will be changed to accommodate hydrophobic mismatch between the protein and micelle (*c* → *c*′), and the semi-axes *a* and *b* are expanded or shrunk (*a* → *a*′ and *b* → *b*′), keeping the volume of the surfactant hydrophobic core region constant. *m*_1_ and *m*_2_ in eq 1 and *s* in eq 5 may also change (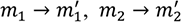, and *s* → *s*′). Hereafter, the prime indicates a parameter in the presence of membrane proteins. To estimate these variables we propose a general scheme below, in which empirical parameters were derived from all-atom MD simulations of protein-micelle complexes as well as pure micelles.

**Figure 2.**
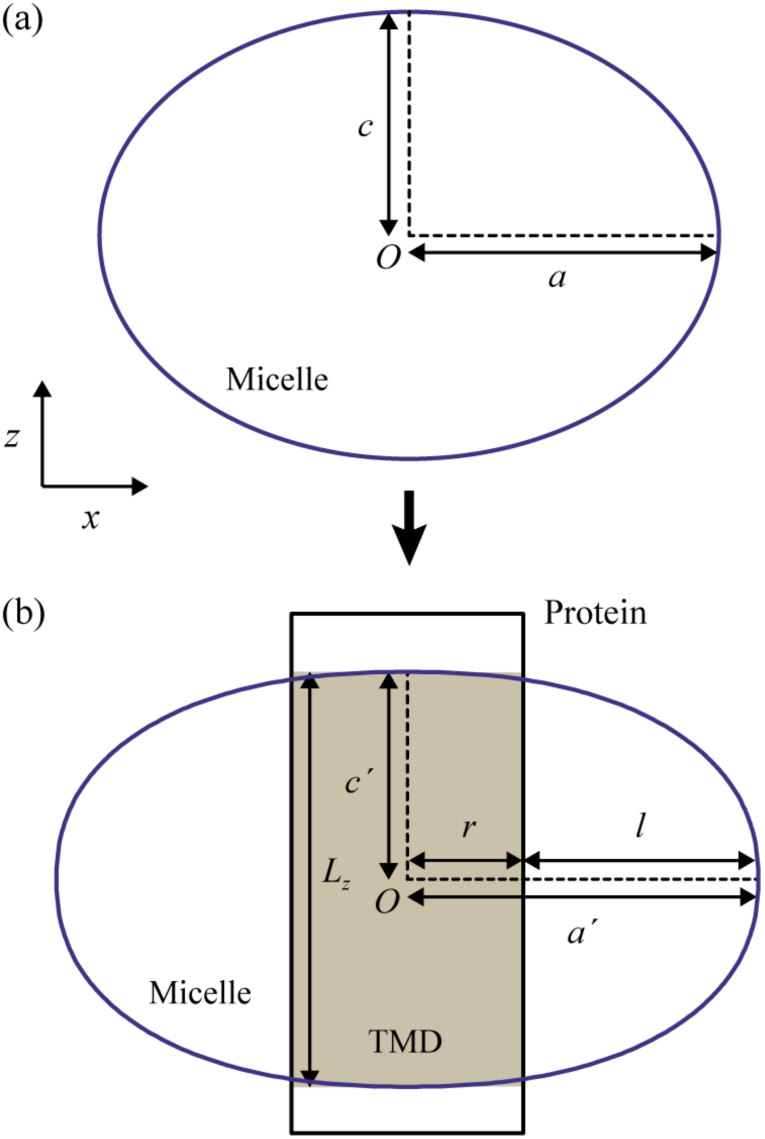
Changes in the micelle size and shape upon protein insertion. (a) Pure micelle and (b) protein-micelle complex. The blue lines stand for the boundary between water and surfactant hydrocarbon groups, and the shaded region colored by brown is the transmembrane domain (TMD) of the embedded protein.

### All-atom MD simulations of pure micelles

We carried out all-atom MD simulations of pure micelles in explicit solvent. We chose decyl-phosphocholine (Fos10), dodecyl-phosphocholine (DPC or Fos12), tetradecyl-phosphocholine (Fos14), sodium dodecyl sulfate (SDS), lauryldimethylamine *N*-oxide (LDAO), and dihexanoyl-phosphatidylcholine (DHPC) to examine how parameters *a, b, c, m*_1_, *m*_2_, and *s* depend on surfactant headgroup type and hydrocarbon length. We modeled the micelles with various aggregation numbers of surfactant by using the CHARMM-GUI *micelle builder* (Table SI),^26^ and carried out a 100-ns MD simulation in the *NPT* ensemble at 300 K and 1 atm for each system (for detailed simulation conditions, see Supporting Information).

As reported previously,^67,68^ micelle structure was highly dynamic, and its instantaneous shapes were irregular. We found that the averaged shape was an ordinary ellipsoid, especially when the micelle was composed of a typical aggregation number of surfactant. Figure 3 shows the mass density profile of 60DPC in the *X* and *Y* dimensions. By fitting a super-ellipsoid function to the three-dimensional (3D) density profile, we obtained *a* = 19.2 Å, *b* = 16.5 Å, *c* = 14.3 Å, *m*_1_ = 0.98, *m*_2_ = 1.00, and *s* = 0.50. In most other cases, *m*_2_ was close to 1.0 (see Table SI). *m*_1_ slightly decreased as *N*_surf_ increased, presumably because the flat region increased as in bicelles. Both *a* and *b* increased as *N*_surf_ increased, while *c* did not seem to exceed a certain length (e.g., 15.1 Å for DPC micelles). The ratio of *a* and *b* was 1.15-1.21 in most cases. The moments of inertia of the ellipsoid calculated from *a, b*, and *c* were comparable to those reported in recent simulation studies.^58^ *s* was slightly dependent on the surfactant headgroup type, where *s* = ∼0.5 for DPC, Fos12, and Fos14 and ∼0.4 for SDS, LDAO, and DHPC. The volume per surfactant (*V*_surf_) was approximately proportional to the number of hydrocarbon atoms, and determined as *V*_surf_ = 322 Å^3^ for DPC, SDS, and LDAO, 268 Å^3^ for Fos10, 379 Å^3^ for Fos14, and 488 Å^3^ for DHPC.

**Figure 3.**
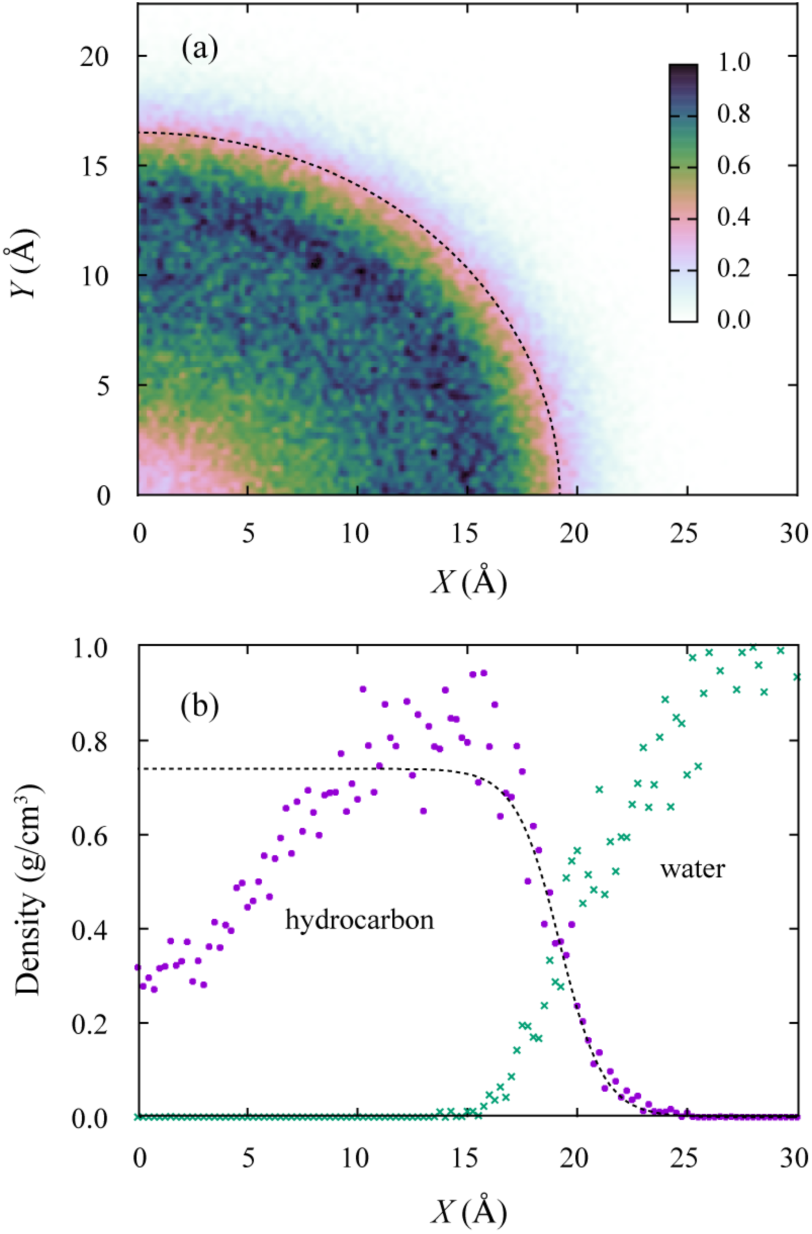
Mass density profiles of the hydrocarbon group in the pure 60DPC micelle. Dashed line represents the best fitting super-ellipsoid. (a) 2D density profile at the *XY* plane. The unit of the density is g/cm^3^. (b) 1D density profile at *y* = 0 and *z* = 0.

### All-atom MD simulations of protein-micelle complexes

All-atom MD simulations were again carried out for the twelve selected membrane-proteins (Hemagglutinin fusion peptide, Integrin β3, Glycophorin A, TMEM14A, AchR β2, BcTSPO, semiSWEET, D3 receptor, KcsA, OmpX, OmpA, and TtoA) in micelles with various aggregation numbers of surfactant (see Figure 4a). We employed 36 systems in total (see Table SII), and performed a 100-200 ns run for each in the *NPT* ensemble at 300 K and 1 atm (for detailed simulation conditions, see Supporting Information). Hereafter, we call these systems “test sets”, and the results were repeatedly used in this study for comparison with the MD simulations in the IMIC model. We analyzed how protein size and surfactant aggregation number affect parameters 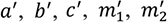, and *s*′ based on the mass density profile of the surfactant hydrocarbon group.

**Figure 4.**
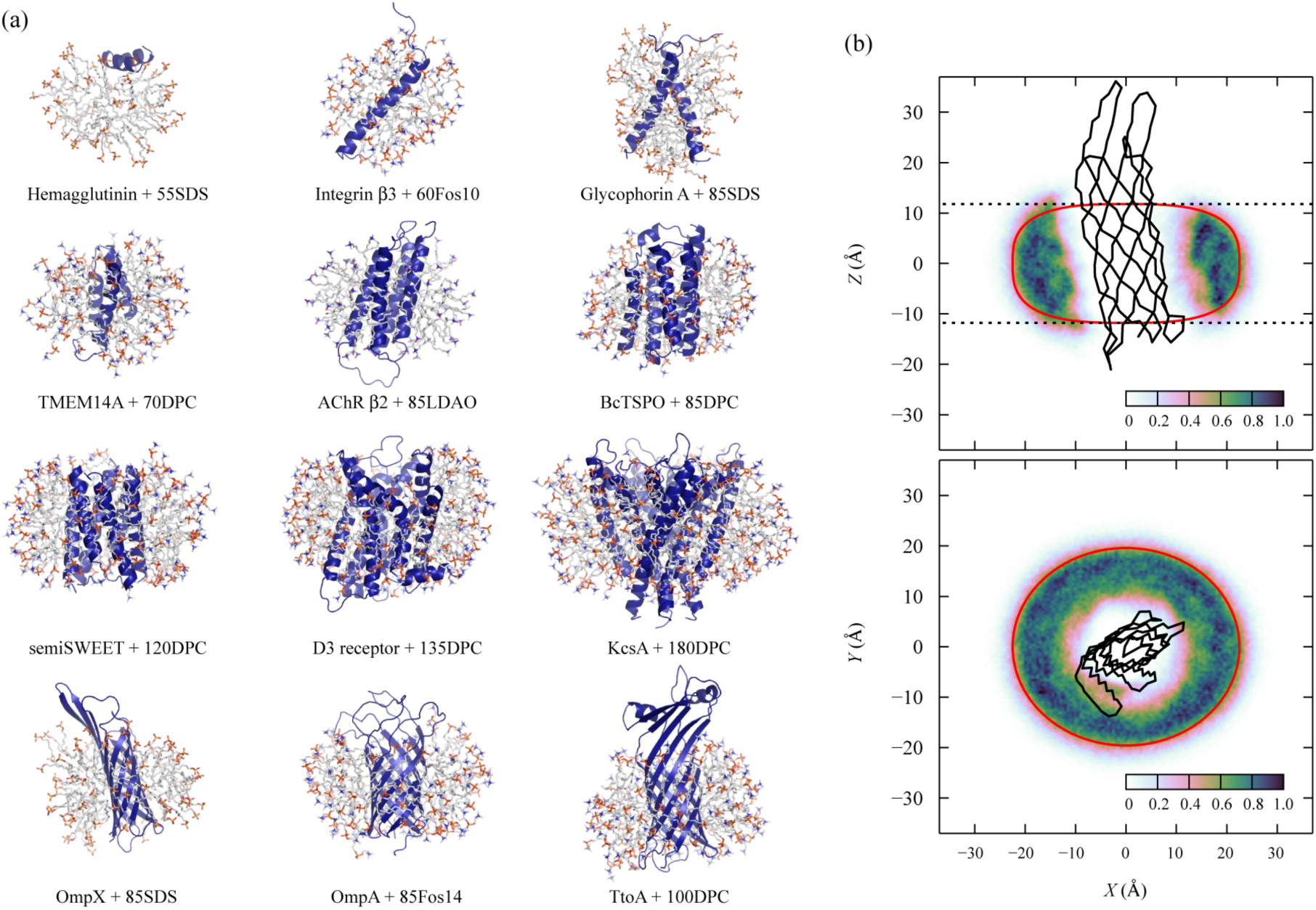
Structure of protein-micelle complexes in explicit solvent. (a) Snapshots of the selected test systems. (b) Mass density profile of the surfactant hydrocarbon group in the OmpX-60DPC complex calculated from the all-atom MD simulation with explicit solvent. The unit of the density is g/cm^3^. Black solid line is the averaged structure of OmpX in the reoriented micelle, dashed line stands for the membrane thickness in the OPM database, and red line is the super-ellipsoid predicted from eqs 7–14.

Figure 4b illustrates the density profile obtained in the OmpX-60DPC complex, where the black solid line is the averaged backbone structure of OmpX in the reoriented micelle. As expected, the whole micelle structure showed an ellipsoidal shape, and the protein was located near the micelle center. Interestingly, the hydrophobic core width of the micelle near the protein along the *Z*-axis was close to the membrane thickness in the OPM database (23.6 Å) (dotted lines in Figure 4b). The averaged micelle thickness in the *XY*-plane was 11.7 Å. Similar results were obtained in the other test systems (Figure S2).

We calculated the super-ellipsoid that fits the micelle-water interface of the mass density profile. In the case of OmpX-60DPC, we obtained *a*′ = 23.4 Å, *b*′ = 20.3 Å, *c*′ = 11.8 Å, 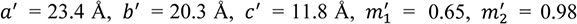, and *s*′ = 0.51. Among the 36 systems, we found some common features and obvious tendencies in the parameters (see Table SIII): 1) *a*′*/b*′ = ∼1.15, 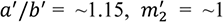, and *s*′ = 0.35-0.5 in most cases. 2) 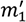 decreases as the protein size in the *X-Y* dimension increases or hydrophobic thickness decreases. (3) Surfactant headgroup type can affect *s*, where SDS showed a smaller *s*′ than DPC, Fos10, and Fos14. (4) In the case of membrane bound peptides, the micelle size is approximately the same as that of the pure micelle. (5) In the case of large membrane proteins, micelle shape can be strongly deformed due to the protein shape (Figure S2), and thus, 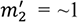 may not be applicable to large proteins.

### General scheme to estimate the micelle size and shape

Based on the above results, we propose a scheme to estimate the micelle size and shape in the presence of membrane proteins without performing all-atom MD simulations. As shown in Figure 2, the total volume of the protein-surfactant complex (*V*_complex_), excluding the extra-micellar region, can be described as a sum of the volumes of the TM domain (*V*_TMD_) and surrounding surfactants:

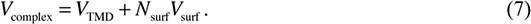

*N*_surf_ is the aggregation number of surfactant, and *V*_surf_ is the volume per surfactant molecule. *V*_complex_ can be calculated by

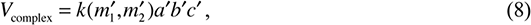

Where

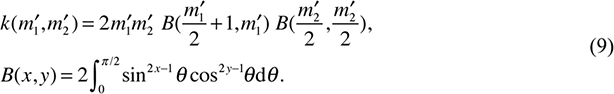

*B*(*x,y*) is the so called beta function. *V*_surf_ can be estimated from the volume of a pure micelle composed of *N*_surf_ surfactants (Figure 2a), given by

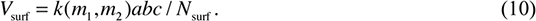

Now, we approximate the embedded protein as a cylindrical shape with hydrophobic (or TMD) thickness *L*_z_ and effective radius *r* (see Figure 2b). To fully solvate TMD with the micelle, *c*′ is set to half of the hydrophobic thickness of the protein:

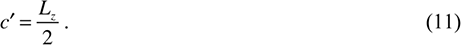

The effective radius of the protein is calculated by

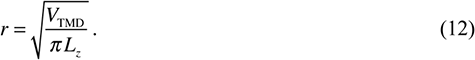

Micelle width *l* on the *XY*-plane can be roughly estimated by assuming that the protein is surrounded by surfactants whose cross-section in a plane perpendicular to the protein surface is semi-ellipsoid and its semi-axes are *l* and *L*_z_/2. Based on the Pappus-Guldinus theorem about *Z*-rotation, *l* is readily calculated as

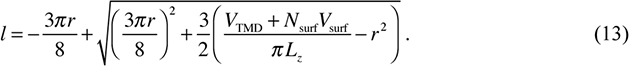

For the estimation of 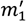, we introduce an empirical formula:

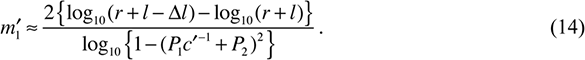

where *P*_1_ and *P*_2_ are the parameters that decide the vertical width of the super-ellipsoid at *x* = *r* + *l* − Δ*l* and *y* = 0. We use Δ*l* = 2 Å, *P*_1_ = 2.761 and *P*_2_ = 0.259, which were derived from a micelle surface curvature calculated from the all-atom MD simulations of protein-micelle complexes. Note that if the estimated 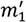 exceeds the lower limit 0 or upper limit 1, it must be fixed to the limit.

Finally, *a*′, *b*′, and 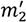 are estimated empirically. As described above, our MD simulations for various protein-micelle complexes demonstrated that *a*′*/b*′ = ∼1.15 and 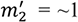 are applicable to many cases, especially when the protein is adequately solvated with surfactant. Accordingly, 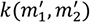 is calculated with eq 9, and thereby, *a*′ and *b*′ are obtained with eq 8.

### Practical estimation of the micelle size and shape

Once *N*_surf_, *V*_surf_, *L*_*z*_, and *V*_TMD_ are decided for a target system, we can estimate 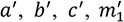, and 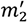 from eqs 7-14. In the case of OmpX-60DPC, we specify *N*_surf_ = 60, *V*_surf_ = 322 Å^3^, *L*_*z*_ = 23.6 Å, and *V*_TMD_ = 10,801 Å^3^, where TMD was defined based on the OPM database,^69^ and *V*_TMD_ was calculated from the heavy atoms in TMD using the 3V program.^70^ We obtain *a*′ = 22.5 Å, *b*′ = 19.6 Å, *c*′ = 11.8 Å, 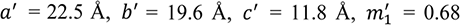, and 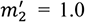, whose shape is illustrated as the red line in Figure 4b. Our estimation showed good agreement with the density profile in the all-atom MD simulation. Similar results were obtained in the other protein-micelle complexes (Table SIV and Figure S2). Among the 36 systems, the standard error of the estimate for *a*′, *b*′, and 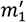 was only 1.5 Å, 1.3 Å, and 0.08, respectively (Figure S3). These results indicate that our scheme is entirely appropriate for estimating the micelle size and shape in the presence of membrane proteins with a given aggregation number of surfactant molecules. As for *s*′, we use 0.35 for SDS and 0.45 for DPC, Fos10, and Fos14. The estimated parameters can now be used for simulating membrane proteins in the IMIC model.

## RESULTS

### Comparison between the IMIC and all-atom models

To validate the IMIC model, we carried out MD simulations for the 36 test systems (see Table SIV), and compared structural and dynamic properties of membrane proteins with those in the all-atom MD simulations. To prepare the initial structure in the IMIC model, the X-ray crystal or NMR structure was placed near the micelle center with the same orientation as in OPM, and rotated about the *Z*-axis so that the PC1 and PC2 axes of the *X,Y*-coordinates of the TM atoms were aligned with the *X,Y*-axes. We conducted a short energy minimization, followed by a 100-200-ns MD simulation at 300 K using the Langevin thermostat. The equations of motion were integrated with the leapfrog method with a time step of 2 fs, where the SHAKE algorithm was used for bond constraint. We used a cutoff distance of 25 Å for the non-bonded interactions. To prevent the embedded proteins from large lateral shifting, we applied a weak restraint on the distance from the center of mass of the TM domain to the *Z*-axis using a flat-bottom restraint potential, where we used 1.5 Å for the switching distance between flat and harmonic functions, and 1.0 kcal/mol·Å^2^ for the force constant.

Here, we focus on OmpX-60DPC. Figure 5a shows the snapshot of OmpX at 200 ns in the IMIC_60DPC_ model. The orientation of OmpX with respect to the *Z*-axis was similar to that in the all-atom MD simulation (compare with Figure 4b). Figure 5b shows the time courses of the root-mean-square deviation (RMSD) for the all Cα atoms with respect to the X-ray crystal structure (PDB entry: 1QJ8). In both IMIC and all-atom models, the RMSD was less than 3 Å over 200 ns, indicating that OmpX was stable in the 60DPC micelle, and this is also consistent with experiment.^71^ Figure 5c shows the root-mean-square fluctuation (RMSF) of the Cα atoms. The RMSF in the IMIC model is in good agreement with that in the all-atom model. The averaged difference between the two RMSF profiles, which we define as 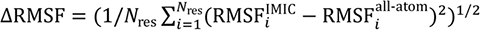 excluding the N- and C-terminal two residues, was 0.134 Å in the transmembrane secondary structure regions, and 0.639 Å in the other regions.

**Figure 5.**
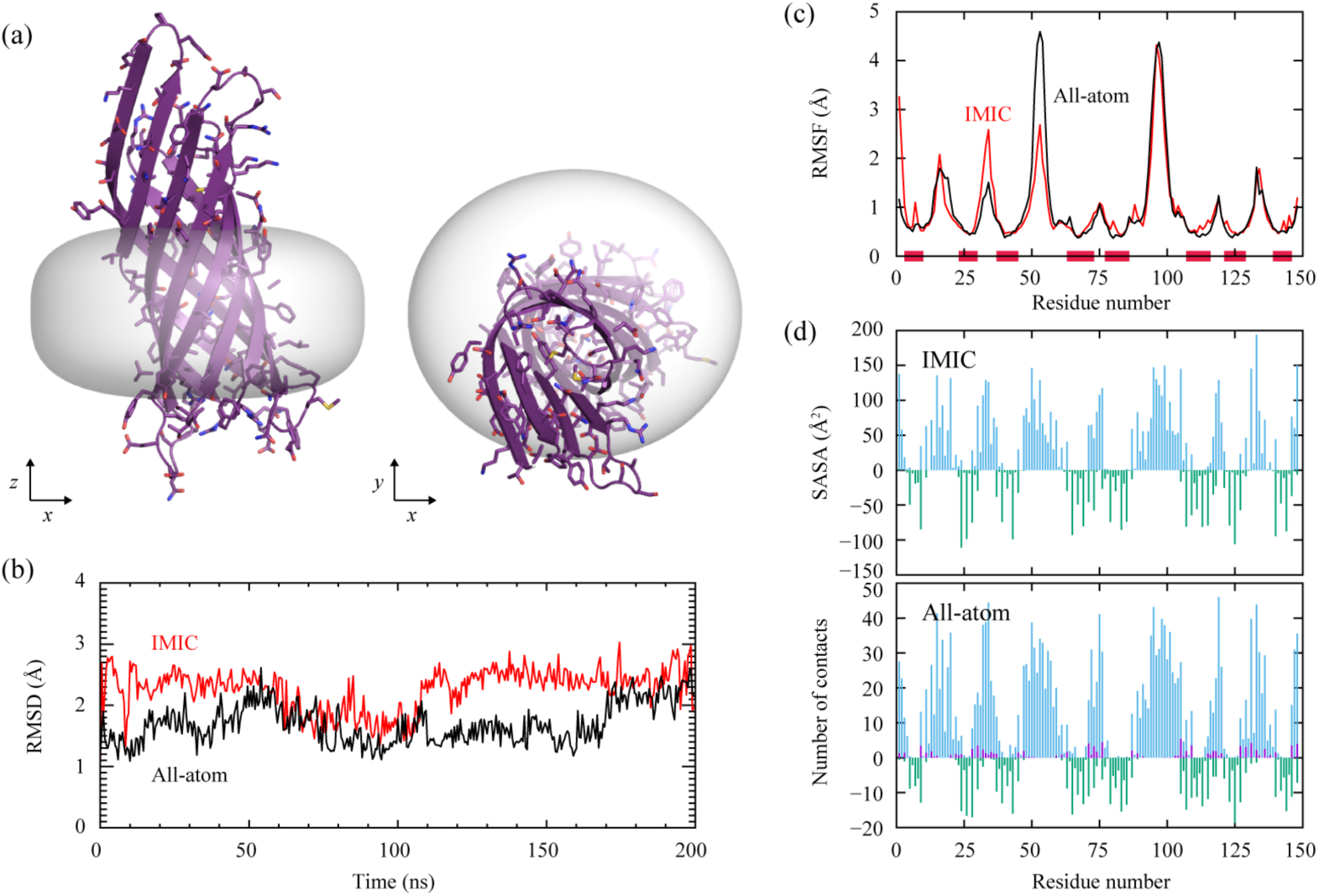
Comparison of the structural and dynamic properties of OmpX in 60DPC in the IMIC and all-atom models. (a) Snapshot of OmpX in the IMIC model at 200 ns. The micelle hydrophobic core region is represented as surface model. (b) RMSD for the all Cα atoms with respect to the X-ray structure. (c) RMSF of the Cα atoms. The horizontal red line denotes the transmembrane β-strand region derived from the OPM database. (d) Solvent accessibility (SA) of each residue in the IMIC (upper panel) and all-atom (lower panel) models. Blue, purple, and green lines are for water, surfactant head group, and surfactant hydrocarbon group, respectively. Note that the SA for the surfactant hydrocarbon group is illustrated as the negative value for the sake of clear visualization.

We also analyzed the degree of burial of OmpX in the micelle by calculating the solvent accessibility (SA) of each residue. In the IMIC model, we calculated the averaged solvent accessible surface area using a Monte Carlo integration scheme, where the water-SA and surfactant-SA were calculated individually according to the position of the probe particle (1.4 Å in radius). In the all-atom model, we calculated the averaged number of solvent atoms around the residues using a cutoff distance of 4 Å. Note that the contacts with the surfactant head groups were included in the water-SA. Figure 5d compares the SA profiles in the IMIC (upper panel) and all-atom models (lower panel). Both two models showed similar profiles, where the β-strands and loops were mainly solvated with surfactant hydrocarbon groups and water, respectively. Correlation coefficients (c.c.) between the two profiles were 0.93 for water-SA, and 0.95 for surfactant-SA, demonstrating that the IMIC model can reasonably mimic a micelle environment.

Similar results were obtained in most other cases of the test sets (see Figure S4a). In both IMIC and all-atom models, the RMSD for the Cα atoms with respect to the initial structure was 4-6 Å in helix bundle proteins with 2-5 TM helices (e.g., GpA, TMEM14A, and AChR β3), while 2-3 Å in the bigger or β-barrel proteins (e.g., D3 receptor, KcsA, and TtoA). *Δ*RMSF for the secondary structure region in TMD was typically less than 0.5 Å, and large *Δ*RMSF (> 1.5 Å) was only observed in the non-secondary structure region of AChR β2 and OmpA, both of which have large extra-micelle loops. The solvent accessibility showed high c.c. (> 0.8) in most cases except for GpA-100DPC (c.c. = 0.48 in surfactant-SA) and TMEM14A-45DPC (c.c = 0.41 in surfactant-SA). This is presumably because 100DPC is excessive for GpA, and 45DPC insufficient for TMEM14A, which might cause a different solvent exposure of proteins in the IMIC and all-atom models. Note that in these cases c.c. is still positive and the profiles were similar to each other. Overall, the IMIC model can reasonably reproduce the all-atom model.

### Glycophorin A in the IMIC, IMM1, and all-atom models

We further focus on glycophorin A (GpA). Experiments have demonstrated that GpA forms an identical conformation in the DPC micelles,^72^ DMPC:DHPC bicelles,^21^ and monoolein lipidic cube phase (LCP).^73^ Among these conditions, the bicelles and LCP are closer to a lipid bilayer environment. An aggregation number of ∼80 for C_12_ surfactants around GpA was also determined.^28^ We simulated GpA in the IMIC, IMM1, and all-atom explicit micelle/membrane models, and compared them with experiments. We employed 60DPC, 85DPC, and 100DPC micelles, and DMPC, DOPC, and POPC bilayers (for detailed simulation conditions, see Supporting Information).

Figure 6a shows a representative snapshot of GpA in the IMIC_85DPC_ model. There were tight G_79_xxxG_83_ motif interactions between monomers over 200 ns. The averaged inter-helical distance *d*_HH_ was 6.1 Å, and the averaged inter-helical crossing angle *θ* was −43.0°. These results were in agreement with those in the all-atom 85DPC model (Figure 6b) (*d*_HH_ = 5.8 Å and *θ* = −44.1°) as well as experiments in DPC micelles (*d*_HH_ = 6.3 Å and *θ* = −41.2° in PDB: 2KPE).^21^ The obtained structures in the DPC micelles were similar to those in the DMPC bilayers (Figures 6c and 6d and Table I), again consistent with experiments.^21^ In both micelles and bilayers, *θ* slightly decreased as the micelle size or membrane thickness increased (see Table II), which is mainly due to hydrophobic matching between GpA and micelles or bilayers. The effective energy *W* in the IMIC model was lower than that in the IMM1 model, where we obtained *W* = 209.1 kcal/mol in IMIC_85DPC_, while *W* = 213.8 kcal/mol in IMM1_DMPC_ (see also Table SV). This is presumably because GpA can easily rotate inside the micelle to accommodate the hydrophobic matching, resulting in a less strained conformation compared to that in the lipid bilayers.

**Table I.**
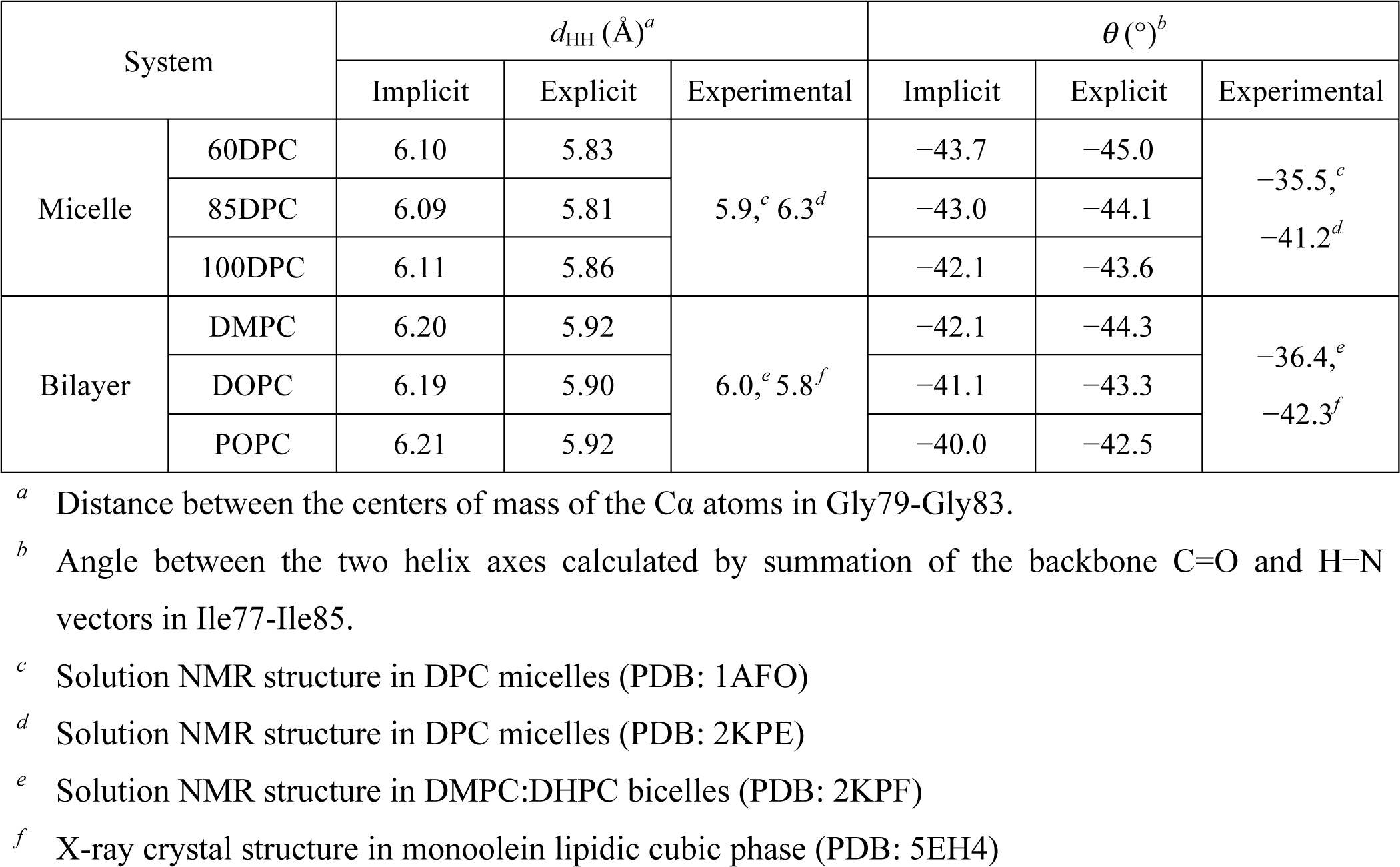
Comparison of the inter-helical distance *d*_HH_ and crossing angle *θ* of the glycophorin A dimer in micelles and lipid bilayers using the implicit (IMIC/IMM1) and all-atom explicit solvent/micelle or membrane models.

**Figure 6.**
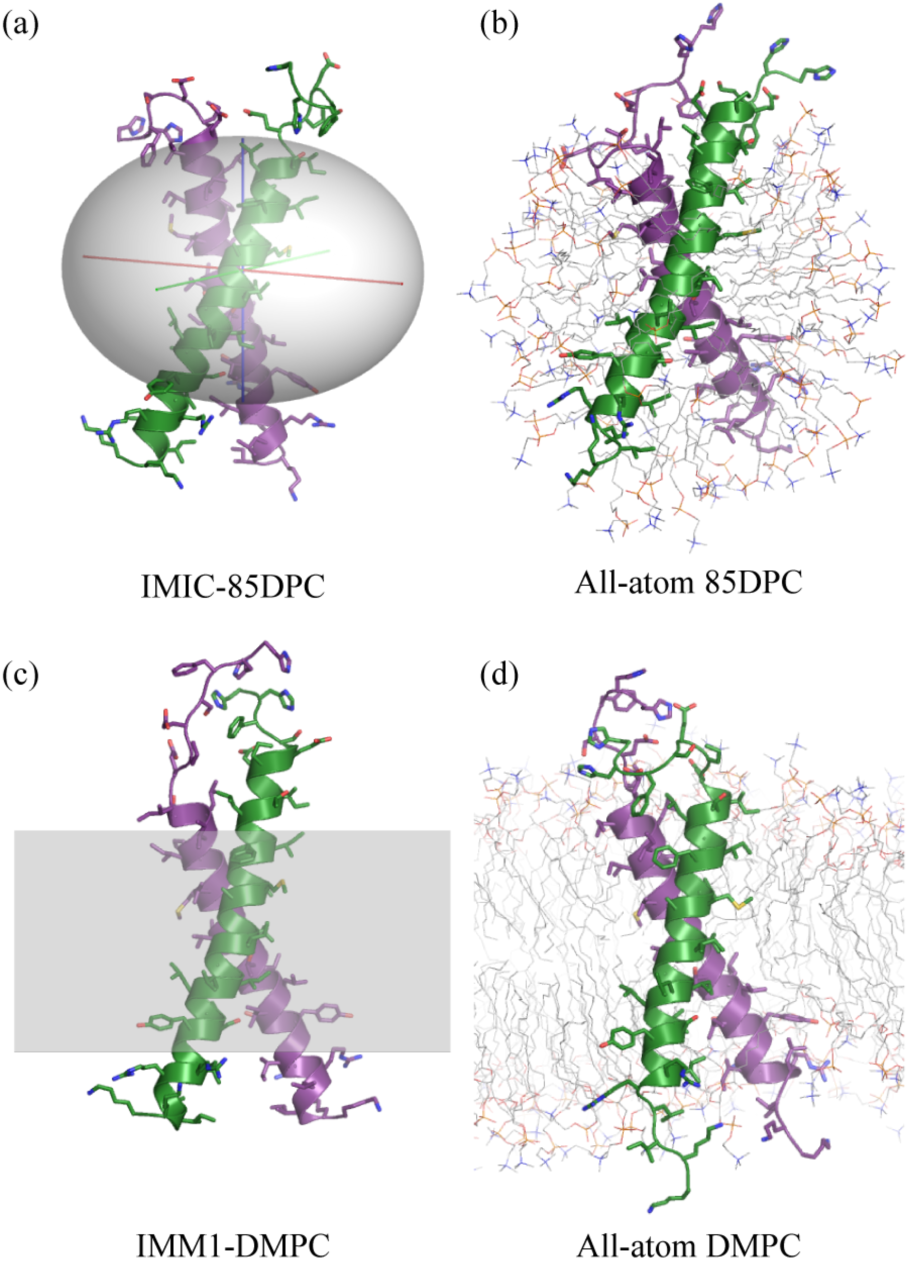
Representative snapshots of GpA in 85DPC micelles and DMPC bilayers. The red, green, and blue lines in the ellipsoid show *X, Y*, and *Z*-axes, respectively.

### Curved surface effects on protein dynamics

To examine curved-surface effects of micelles on membrane protein structure, we simulated the HIV-1 envelope glycoprotein 41 (gp41). This protein has an important role in membrane fusion between the viral envelope and cellular membranes.^74^ It has a long C-terminal cytoplasmic tail, consisting of three amphipathic α-helical portions: LLP (lentiviral lytic peptide)-1, 2, and 3. Fluorescence and infrared spectroscopy demonstrated that LLP-3 tightly binds to lipid bilayers, and the helix axis is nearly parallel to the membrane plane (∼70°).^75^ Moreover, NMR experiments suggested that the α-helix of LLP-3 is almost straight on the DHPC:DMPC bicelle surface, while it is slightly curved on the DHPC micelle, indicating a curved-surface effect of micelles.^12^

We carried out MD simulations of LLP-3 using the IMIC and IMM1 models, and compared the structures in the two environments. We employed a 30DHPC micelle and DMPC lipid bilayer, and performed long MD runs (500 ns in IMIC and 150 ns in IMM1 at 300 K) to search the optimal position and orientation of the peptide on the micelle/bilayer surface, starting from the ideal α-helical conformation. Figure 7a illustrates a representative structure of LLP3 in the IMIC model, whose probability of spatial position and orientation was highest according to Euler angle analysis (Figure S5). LLP-3 adopted an α-helix on the micelle surface, and the side-chains of Trp5, Leu9, Trp12, and Leu16 were fully buried in the hydrophobic core region. Similarly, in the IMM1 model, LLP-3 bound to the bilayer surface (Figure 7b), and the averaged tilt angle was ∼86°, which is reasonably consistent with the experimental data (∼70°).^75^

**Figure 7.**
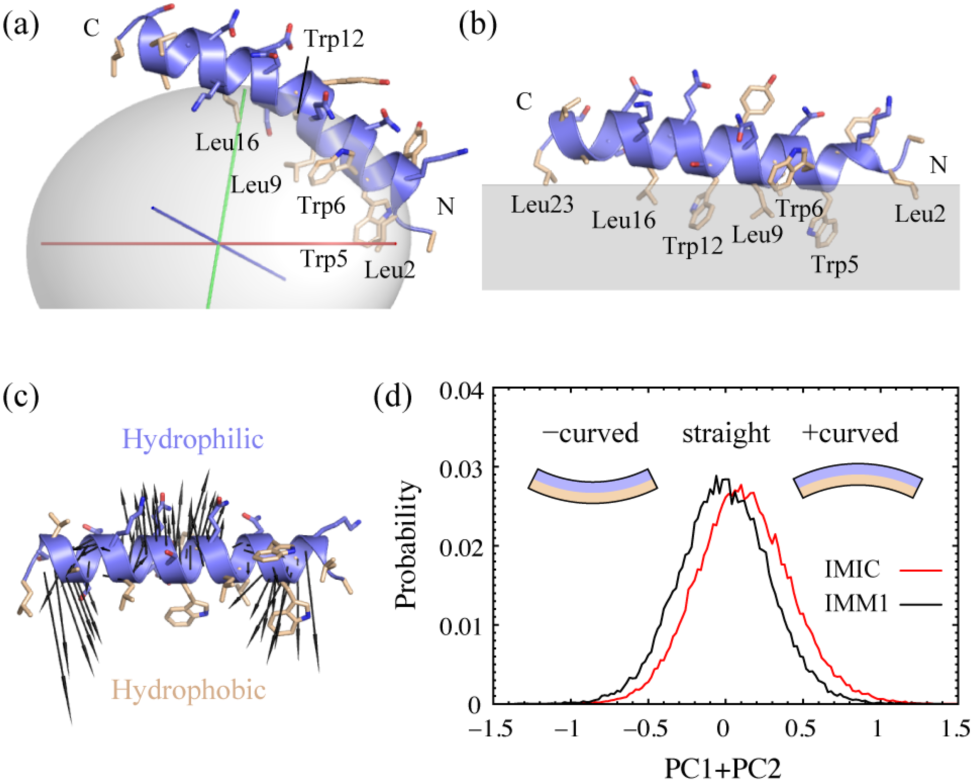
Structure and dynamics of the LLP-3 peptide on the IMIC and IMM1 model surfaces. (a) Representative snapshot in the IMIC_30DHPC_ model, and (b) representative snapshot in the IMM1_DMPC_ model. (c) Helix bending motion obtained from the principal component analysis for the IMM1 simulations, where the PC1 and PC2 vectors were summed. (d) Probability distribution of the projection of the trajectories on the (PC1+PC2) vector. The inset illustrates a curved α-helix, where the blue and orange colors stand for the hydrophilic and hydrophobic sides, respectively.

To investigate dynamic properties of LLP-3 on the micelle and bilayer surfaces, we carried out principal component analysis (PCA) for the MD trajectories. In both IMIC and IMM1 models, the first and second principal components (PC1 and PC2) were bending motions of the α-helix, which were perpendicular to each other. These two modes contributed about 48% to the total fluctuations. We found that the sum of the PC1 and PC2 vectors gave a bending motion from the hydrophilic side of the helix towards the hydrophobic side, and vice versa (Figure 7c), and the direction appeared to be perpendicular to the micelle/bilayer surface. We projected the trajectories onto the (PC1+PC2) vector obtained from the IMM1 simulation, and calculated their distributions (Figure 7d). In the IMM1 model we obtained a Gaussian distribution with zero mean, while in the IMIC model it was shifted towards positive values. Here, the averaged bending angle of the helix at and around 0.0, 0.2, 0.5, and 1.0 on the (PC1+PC2) axis was 175, 173, 169, and 162°, respectively. These results indicate that the structural fluctuation of LLP3 on the micelle surface is biased to the “+curved” conformation, in which the hydrophilic side of the α-helix is relatively extended (insets of Figure 7d), consistent with NMR experiments.^12^ The IMIC model faithfully induces the curved α-helix conformation on the micelle surface.

### Compact hydrophobic-core effects on protein conformation

We simulated the bacteriophage M13 major coat protein gp8. This is a 50-residue protein that assembles around the phage particle to encapsulate the circular single-stranded DNA, and has an important role in interacting with the host membrane in the infection process.^76^ Site-directed labeling (SDL) experiments revealed that gp8 adopts a continuous α-helix (I-shape) in fully hydrated DOPC:DOPG vesicles.^77^ Solid-state NMR showed a helix-turn-helix conformation (L-shape), consisting of the TM and juxtamembrane (JM) helices, in oriented POPC:POPG lipid bilayers.^78^ Solution NMR showed various conformations in SDS micelles, including not only I- and L-shapes but also U-shape.^79^ These experiments suggest that I- and L-shapes can commonly exist in micelle and bilayer environments, while U-shape is an artifact in the membrane mimetic environments.

To examine these characteristics, we performed umbrella sampling of gp8 in the IMIC_60SDS_ and IMM1_POPC_ models with the reaction coordinates of kink angle between the TM and JM helices. The amino-acid sequence of gp8 is shown in Figure 8a. We employed 18 window potentials, where the angle between the centers of Cα atoms of Ala9-Phe11, Thr19-Tyr21, and Lys44-Thr46 was restrained at 10° intervals from 10 to 180° with the force constant of 40 kcal/mol·rad^2^ for each window. We used U-shape (Model 8 in PDB entry: 2CPS) for the initial structure, and performed a 50-ns equilibration, followed by 50-ns production run for each window. The free energy profile was computed with the weighted histogram analysis method (WHAM).^80,81^

**Figure 8.**
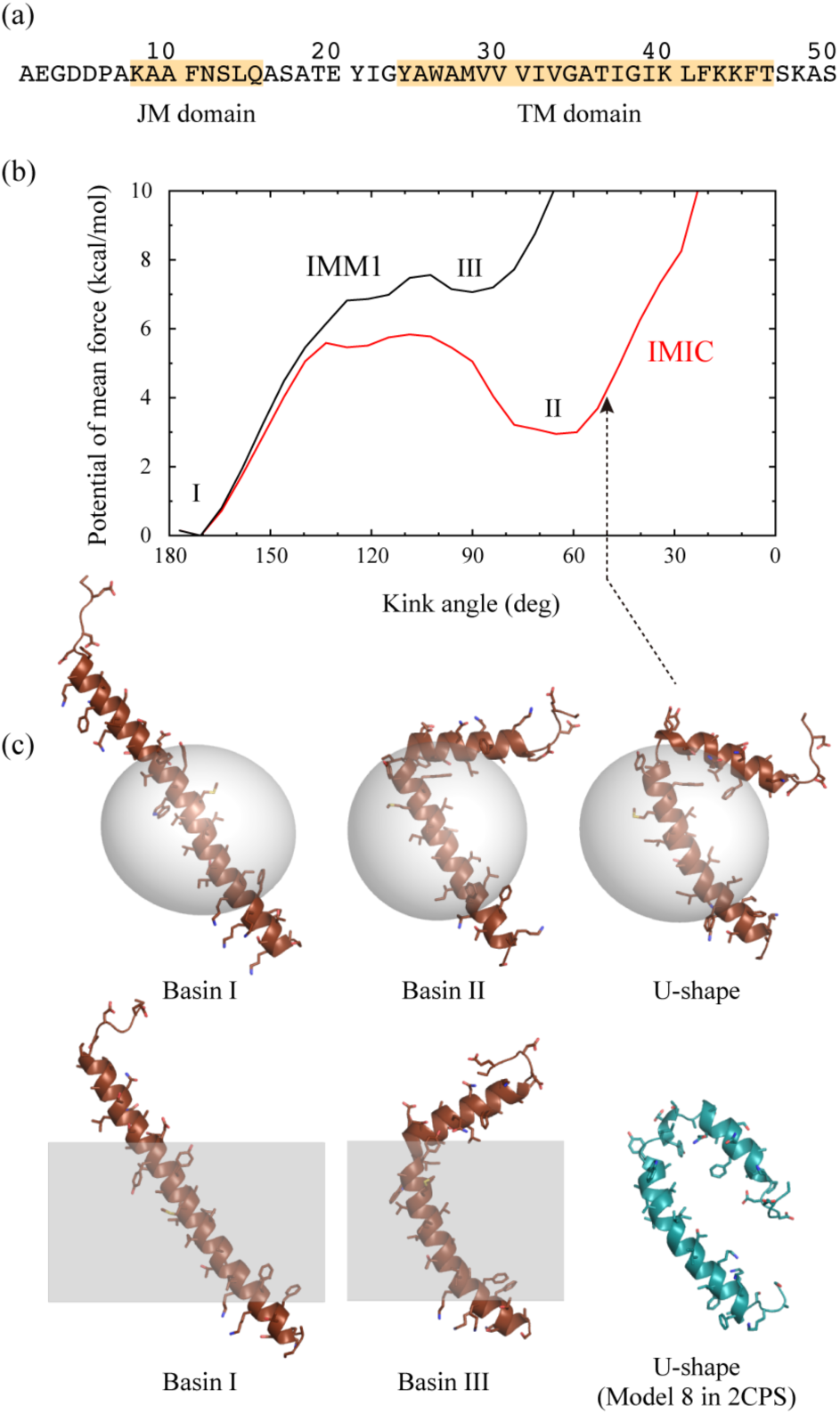
Umbrella sampling simulations of the bacteriophage M13 major coat protein gp8 in the IMIC and IMM1 models. (a) Amino-acid sequence of gp8. (b) Potential of mean force at 298.15 K as the function of the kink angle between the TM and JM helices. The *X*-axis was reversed for the sake of clarity. (c) Representative structures in Basins I-III.

Figure 8b shows the potential of mean force (PMF) at 298.15 K as the function of the kink angle. In both IMIC and IMM1 models, the global energy minimum state was I-shape, where the kink angle was ∼180° (Basin I). In the IMIC model, there was another deep and wide basin around ∼64° (Basin II), which was mainly composed of L-shape. Free energy difference between I- and L-shapes was 3.0 kcal/mol. Conformational transition between L- and U-shapes (∼51°) may occur due to thermal fluctuation in this basin. In the IMM1 model, L-shape was also found in a local energy minimum at 90° (Basin III), but the free energy difference from I-shape was very large, suggesting that I-shape is predominant in lipid bilayers. As suggested by Vos *et al.*, L-shape in membranes might be specifically stabilized in the oriented lipid bilayers, where the stacked bilayers can press the JM domain into the membrane-water interface.^13^

In the stable L-shape obtained in the IMIC model, most hydrophobic residues of the JM helix penetrated into the micelle surface, while a few hydrophobic residues in the IMM1 model. This is presumably because the TM domain can easily rotate or tilt inside the micelle, compared to the lipid bilayers, to accommodate hydrophobic matching in the TM domain and search optimal interactions between the JM helix and curved micelle surface. U-shape seems to be energetically unfavorable in lipid bilayers, since the N-terminal hydrophilic residues in the JM helix must be deeply inserted into the hydrophobic core region of the membrane. In micelle environments, it can be avoided because of the compact hydrophobic core space of the micelle. We suggest that accommodation of the interactions between the JM helix and micelle/membrane surface can make a structural difference of membrane proteins in different membrane environments.

### Prediction of the transmembrane domain structure of APP dimer

Amyloid precursor protein (APP) is an integral membrane protein that plays an important role in the pathophysiology of Alzheimer’s disease.^82^ It has been suggested that the APP dimer forms different conformations in lipid bilayers and surfactant micelles. Solid-state NMR experiments demonstrated that APP adopts a right-handed Gly-in conformation in mixed DMPC:DMPG lipid bilayers, where GxxxG motif interactions are formed at the interface between the monomers.^18^ On the other hand, solution NMR experiments in DPC micelles showed left-handed Gly-out (PDB entry: 2LOH)^19^ or right-handed Gly-out conformations (PDB entry: 2LZ3),^20^ where GxxxA motif interactions exist at the interface. In computational studies, Miyashita *et al.* predicted Gly-in by using the REMD method^83^ with the IMM1-C19 model.^84^ Dominguez *et al.* performed multi-scale MD simulations combining coarse-grained and all-atom models, and suggested that Gly-in is stable in thick lipid bilayers, while Gly-out in micelles or thin lipid bilayers.^85,86^ They also reported a Gly-side conformation, in which the two helices adapt a parallel orientation to form a zipper-like interface.

To examine whether the IMIC and IMM1 models can reproduce these characteristics, we predicted the transmembrane domain structures of the APP dimer using the REMD method, and compared them with the NMR structures. We employed 60DPC and 85DPC micelles in IMIC, and POPC lipid bilayers with the membrane thickness of 28.8 and 32.0 Å in IMM1. In the initial structure, the two helices were perpendicular to the *XY*-plane, and each helix was rotated around its helical axis at 90° intervals in each replica, resulting in 16 individual conformations. We performed a 100-ns REMD simulation with 16 replicas (1.6 µs in total) for each system. The temperatures were distributed exponentially from 287.63 to 521.39 K, and replica exchange was attempted every 2,000 step (1 step = 2 fs) (for detailed simulation conditions, see Supporting Information).

During the REMD simulations, the two helices were associated and dissociated repeatedly. Both right-handed and left-handed configurations were sampled, where the right-handed ones were frequently observed (see Figure S6). Figure 9a shows the 2D-PMF at 300 K and 447 K along the RMSDs with respect to the left-handed and right-handed Gly-out conformations. The NMR structures of PDB entries: 2LOH and 2LZ3 are shown as blue squares and red circles, respectively, and the structures predicted by the PREDDIMER server^87^ are also plotted as triangles. In all systems, there were mainly two basins at 300 K (Basins I and II). Basin I was composed of Gly-in, which was close to the PREDDIMER structure, and Basin II was Gly-side (see Figure 9b). We found that the averaged effective energy of Gly-in was 2-3 kcal/mol larger than that of Gly-side in IMIC, while the tendency was opposite in IMM1 (see Table SVI). Interestingly, there was another basin (Basin III) at higher temperatures (382.58-521.39 K) in IMIC_60DPC_, IMIC_85DPC_, and IMM1_28.8Å_, while not in IMM1_32.0Å_. Basin III was mainly composed of Gly-out, and close to the NMR structures in DPC micelles.^20^ These results indicate that Gly-out is not so stable compared to Gly-in or Gly-side, but has a possibility to exist in micelles or thinner lipid bilayers.

**Figure 9.**
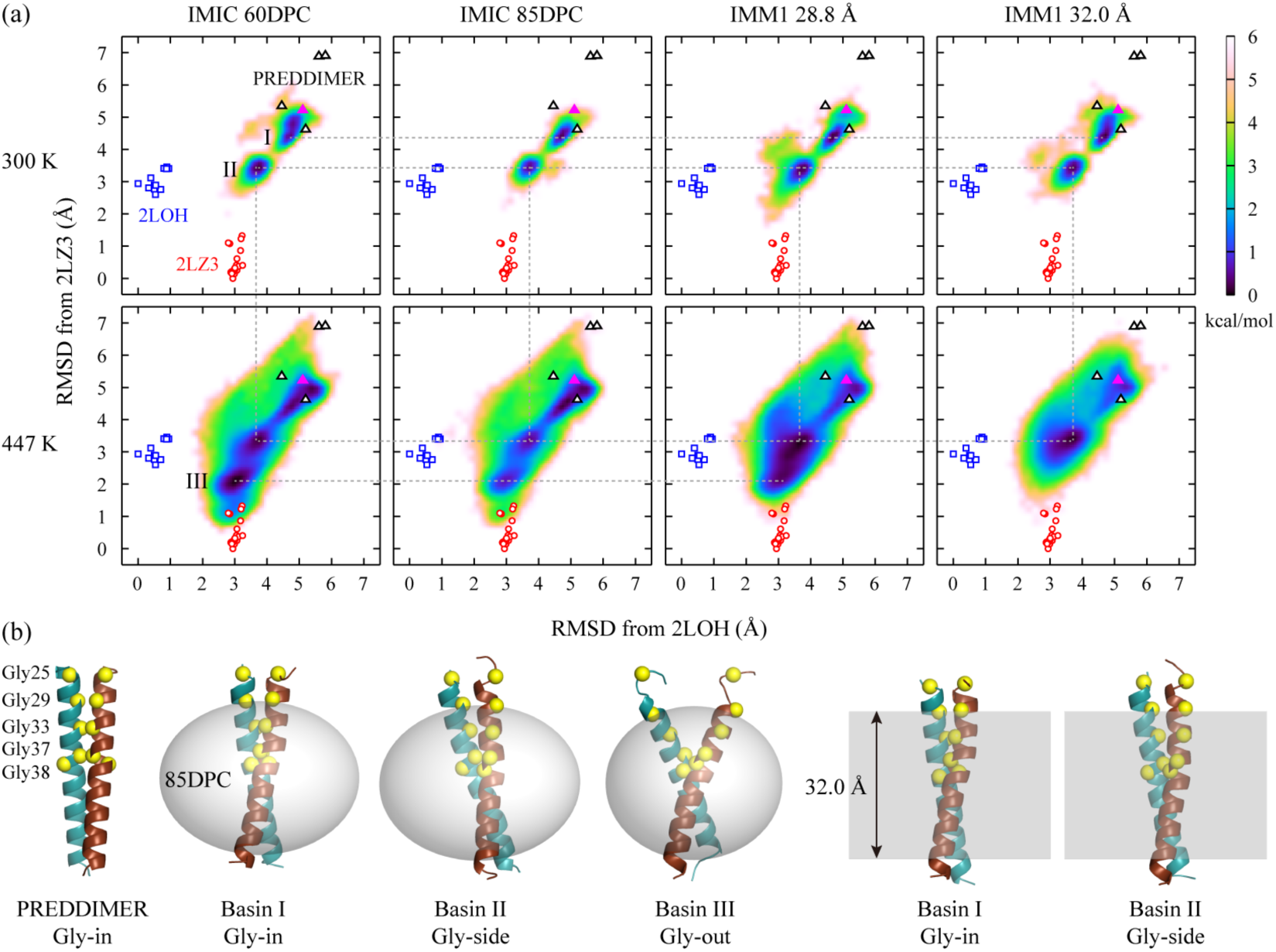
REMD simulations of the APP dimer in the IMIC and IMM1 models. (a) 2D-PMF along the RMSDs with respect to the left-handed Gly-out (Model 1 in PDB: 2LOH) and right-handed Gly-out (Model 1 in PDB: 2LZ3) at 300 K and 447 K. The RMSDs were computed for the Cα atoms of Gly except for Gly25. The red circles and blue squares are the NMR structures of PDB entries: 2LOH and 2LZ3, respectively. The triangles are the five models predicted with the PREDDIMER server, where the magenta stands for the best model. (b) Representative structures obtained from PREDDIMER (magenta), Basins I-III in IMIC_85DPC_, and Basins I and II in IMM1_32.0Å_. The yellow sphere is the Cα atom of Gly.

The curved surface of the micelle is likely to induce formation of Gly-out, since the dimer can adapt an X-shaped structure due to hydrophobic matching. In fact, the averaged helix-crossing angle of Gly-out in IMIC was larger than those in IMM1 [55.6° in IMIC_85DPC_, 57.2° in IMIC_60DPC_, and 49.4° in IMM1_28.8Å_ (see Table SVI)]. Dominguez *et al.* suggested that water molecules that penetrate deeply into micelles or thin lipid bilayers could stabilize the hydrophilic interface in Gly-out.^86^ Such explicit interactions between water and APP might significantly shift the conformational equilibrium from Gly-in to Gly-out in micelles. Consequently, our simulations using the IMIC and IMM1 models are reasonably consistent with the NMR experiments and previous computational studies.

## DISCUSSION

In this study, we developed a new implicit solvent model, named the IMIC model, which treats micelle environments implicitly to simulate protein-micelle complexes at low computational cost. We extended the lipid bilayer model (IMM1) to the micelle system by simply converting the hydrophobic slab to the ellipsoidal shape. This is based on the assumption that the parameters used for the calculation of the solvation free energy in the lipid bilayer model are directly applicable to a micelle model. This is reasonable because the micelle and lipid bilayer are composed of similar chemical compounds. Defining micelle shape with a super-ellipsoid function enables us to create various-shaped systems such as spherical micelles, rod-like micelles, bicelles, and nano-discs merely by changing a few parameters in the function. Our model will be further extended to the PB- and GB-based implicit micelle models.

Because the model proposed here is based on simple formulae and approximations, there is room for improvement. In the current model, only water and cyclohexane phases are considered. As in the IMM1-GC model,^51^ the Gouy-Chapman theory can be combined with the IMIC model, which can introduce one layer between the two phases to describe electrostatic interactions between surfactant headgroups and amino acids. A multiple-layered model has been also proposed for the GB-based implicit membrane model.^47^ It could also be important to incorporate a dynamic feature like local deformation of the micelle into the model, which could be realized by introducing a deformation energy of the micelle into the Hamiltonian, as in the DHDGB model.^48^

The IMIC model will be useful for structure determination of proteins by NMR techniques, especially when the experiments are conducted in micelles. The structure models are usually calculated from simulated annealing MD with geometrical restraints derived from the NMR spectra. It has been demonstrated that the NMR structure calculation in an implicit solvent model is comparable with that in the explicit solvent model.^88^ In addition, Tian *et al* proposed using the IMM1 model for effective structure determination of membrane proteins.^89^ If the target proteins were reconstituted into micelles, it would be reasonable to utilize IMIC rather than IMM1, since the protein structure might be perturbed by the curved surface of the micelles.

Another useful application will be a cryo-EM flexible fitting of membrane proteins.^90^ Cryo-EM is a powerful tool to determine 3D structures of biomolecules with near atomic resolution. In flexible fitting, MD simulation is usually carried out to guide the initial structure towards the target EM density map using a biasing potential. In such protocols, the implicit solvent model is useful to reduce computational cost, and the protein structure can be quickly changed and converged compared to the explicit solvent model.^91^ Experimentally, an ellipsoidal micelle can be clearly observed around the TM domain of membrane proteins in the density maps,^3-6^ enabling us to easily define the hydrophobic core region for the IMIC model. Recently, we have implemented a flexible fitting module into GENESIS,^92^ which is available with not only the EEF1/IMM1/IMIC models but also enhanced sampling algorithms like the replica-exchange MD method.^93^ The integrated methods are capable of realizing reliable structure modeling of membrane proteins in an experimental environment with low computational cost.

## SUMMARY

In this study, we proposed an implicit micelle model (IMIC), which treats micelle environments implicitly. The IMIC model is one of the hydration shell models, and is an extended form of the implicit membrane model IMM1. We introduced a super-ellipsoid function to define the hydrophobic core region of the micelle, and the parameters were derived from all-atom MD simulations of protein-micelle complexes as well as pure micelles. We demonstrate that the structural and dynamic properties of membrane proteins in the IMIC model are similar to those in the explicit solvent model. We compared the conformations of selected membrane proteins (glycophorin A, HIV envelope glycoprotein gp41, bacteriophage M13 major coat protein gp8, and APP dimer) in the IMIC and IMM1 models, and obtained good agreement with NMR experiments. We conclude that the IMIC model faithfully reproduces curved surface effects and compact hydrophobic-core effects of micelles on membrane protein structures, and the combined use of IMIC and IMM1 models enables us to discriminate membrane mimetic artifacts from native structures among experimentally determined NMR structures.

## ACKNOWLEDGMENTS

We would like to thank Drs. Kiyoshi Yagi, Hiraku Oshima, and Donatas Surblys at RIKEN for helpful comments and discussions. This work was supported by JSPS KAKENHI Grant Numbers JP15H05594 and JP26119006, a grant from Innovative Drug Discovery Infrastructure through Functional Control of Biomolecular Systems, Priority Issue 1 in Post-K Supercomputer Development (hp170254), and the RIKEN Pioneering Projects, Integrated Lipidology and Dynamic Structural Biology. MD simulations were partially carried out on HOKUSAI GreatWave and BigWaterFall at RIKEN.

## REFERENCES

(1) Moraes, I.; Evans, G.; Sanchez-Weatherby, J.; Newstead, S.; Stewart, P. D. S. BBA-Biomembranes 2014, 1838, 78–87.

(2) Yeagle, P. L. BBA-Biomembranes 2014, 1838, 1548–1559.

(3) Iadanza, M. G.; Higgins, A. J.; Schiffrin, B.; Calabrese, A. N.; Brockwell, D. J.; Ashcroft, A. E.; Radford, S. E.; Ranson, N. A. Nat. Commun. 2016, 7, 12865

(4) Zhu, S.; Stein, R. A.; Yoshioka, C.; Lee, C.-H.; Goehring, A.; Mchaourab, H. S.; Gouaux, E. Cell 2016, 165, 704–714.

(5) Grieben, M.; Pike, A. C. W.; Shintre, C. A.; Venturi, E.; El-Ajouz, S.; Tessitore, A.; Shrestha, L.; Mukhopadhyay, S.; Mahajan, P.; Chalk, R.; Burgess-Brown, N. A.; Sitsapesan, R.; Huiskonen, J. T.; Carpenter, E. P. Nat. Struct. Mol. Biol. 2017, 24, 114–122.

(6) Zhang, Y.; Sun, B.; Feng, D.; Hu, H.; Chu, M.; Qu, Q.; Tarrasch, J. T.; Li, S.; Kobilka, T. S.; Kobilka, B. K.; Skiniotis, G. Nature 2017, 546, 248–253.

(7) Kang, C.; Li, Q. Curr. Opin. Chem. Biol. 2011, 15, 560–569.

(8) Wang, S.; Ladizhansky, V. Prog. Nucl. Magn. Reson. Spectrosc. 2014, 82, 1–26.

(9) Liang, B.; Tamm, L. K. Nat. Struct. Mol. Biol. 2016, 23, 468–474.

(10) Zhou, H.-X.; Cross, T. A. Annu. Rev. Biophys. 2013, 42, 361–392.

(11) Chipot, C.; Dehez, F.; Schnell, J. R.; Zitzmann, N.; Pebay-Peyroula, E.; Catoire, L. J.; Miroux, B.; Kunji, E. R. S.; Veglia, G.; Cross, T. A.; Schanda, P. Chem. Rev. 2018, 118, 3559–3607.

(12) Chou, J. J.; Kaufman, J. D.; Stahl, S. J.; Wingfield, P. T.; Bax, A. J. Am. Chem. Soc. 2002, 124, 2450–2451.

(13) Vos, W. L.; Nazarov, P. V.; Koehorst, R. B. M.; Spruijt, R. B.; Hemminga, M. A. Trends Biochem. Sci. 2009, 34, 249–255.

(14) Cross, T. A.; Sharma, M.; Yi, M.; Zhou, H.-X. Trends Biochem. Sci. 2011, 36, 117–125.

(15) Oxenoid, K.; Chou, J. J. Proc. Natl. Acad. Sci. U.S.A. 2005, 102, 10870–10875.

(16) Traaseth, N. J.; Verardi, R.; Torgersen, K. D.; Karim, C. B.; Thomas, D. D.; Veglia, G. Proc. Natl. Acad. Sci. U.S.A. 2007, 104, 14676–14681.

(17) Mineev, K. S.; Panova, S. V.; Bocharova, O. V.; Bocharov, E. V.; Arseniev, A. S. Biochemistry 2015, 54, 6295–6298.

(18) Sato, T.; Tang, T. C.; Reubins, G.; Fei, J. Z.; Fujimoto, T.; Kienlen-Campard, P.; Constantinescu, S. N.; Octave, J. N.; Aimoto, S.; Smith, S. O. Proc. Natl. Acad. Sci. U.S.A. 2009, 106, 1421–1426.

(19) Nadezhdin, K. D.; Bocharova, O. V.; Bocharov, E. V.; Arseniev, A. S. FEBS Lett. 2012, 586, 1687–1692.

(20) Chen, W.; Gamache, E.; Rosenman, D. J.; Xie, J.; Lopez, M. M.; Li, Y. M.; Wang, C. Y. Nat. Commun. 2014, 5.

(21) Mineev, K. S.; Bocharov, E. V.; Volynsky, P. E.; Goncharuk, M. V.; Tkach, E. N.; Ermolyuk, Y. S.; Schulga, A. A.; Chupin, V. V.; Maslennikov, I. V.; Efremov, R. G.; Arseniev, A. S. Acta Naturae 2011, 3, 90–98.

(22) Bond, P. J.; Sansom, M. S. P. J. Mol. Biol. 2003, 329, 1035–1053.

(23) Lagüe, P.; Roux, B.; Pastor, R. W. J. Mol. Biol. 2005, 354, 1129–1141.

(24) Rouse, S. L.; Sansom, M. S. P. J. Phys. Chem. B 2015, 119, 764–772.

(25) Dominguez, L.; Meredith, S. C.; Straub, J. E.; Thirumalai, D. J. Am. Chem. Soc. 2014, 136, 854–857.

(26) Cheng, X.; Jo, S.; Lee, H. S.; Klauda, J. B.; Im, W. J. Chem. Inf. Model. 2013, 53, 2171–2180.

(27) Hwang, P. M.; Choy, W.-Y.; Lo, E. I.; Chen, L.; Forman-Kay, J. D.; Raetz, C. R. H.; Privé, G. G.; Bishop, R. E.; Kay, L. E. Proc. Natl. Acad. Sci. U.S.A. 2002, 99, 13560–13565.

(28) Fisher, L. E.; Engelman, D. M.; Sturgis, J. N. Biophys. J. 2003, 85, 3097–3105.

(29) Stanczak, P.; Horst, R.; Serrano, P.; Wüthrich, K. J. Am. Chem. Soc. 2009, 131, 18450–18456.

(30) Chaptal, V.; Delolme, F.; Kilburg, A.; Magnard, S.; Montigny, C.; Picard, M.; Prier, C.; Monticelli, L.; Bornert, O.; Agez, M.; Ravaud, S.; Orelle, C.; Wagner, R.; Jawhari, A.; Broutin, I.; Pebay-Peyroula, E.; Jault, J.-M.; Kaback, H. R.; le Maire, M.; Falson, P. Sci. Rep. 2017, 7, 41751.

(31) Bond, P. J.; Cuthbertson, J. M.; Deol, S. S.; Sansom, M. S. P. J. Am. Chem. Soc. 2004, 126, 15948–15949.

(32) Braun, R.; Engelman, D. M.; Schulten, K. Biophys. J. 2004, 87, 754–763.

(33) Böckmann, R. A.; Caflisch, A. Biophys. J. 2005, 88, 3191–3204.

(34) Lazaridis, T.; Mallik, B.; Chen, Y. J. Phys. Chem. B 2005, 109, 15098–15106.

(35) Bond, P. J.; Sansom, M. S. P. J. Am. Chem. Soc. 2006, 128, 2697–2704.

(36) Jusufi, A.; Hynninen, A.-P.; Panagiotopoulos, A. Z. J. Phys. Chem. B 2008, 112, 13783–13792.

(37) Versace, R. E.; Lazaridis, T. J. Phys. Chem. B 2015, 119, 8037–8047.

(38) Kleinjung, J.; Fraternali, F. Curr. Opin. Struct. Biol. 2014, 25, 126–134.

(39) Honig, B.; Nicholls, A. Science 1995, 268, 1144–1149.

(40) Still, W. C.; Tempczyk, A.; Hawley, R. C.; Hendrickson, T. J. Am. Chem. Soc. 1990, 112, 6127–6129.

(41) Eisenberg, D.; McLachlan, A. D. Nature 1986, 319, 199–203.

(42) Kang, Y. K.; Némethy, G.; Scheraga, H. A. J. Phys. Chem. 1987, 91, 4105–4109.

(43) Lazaridis, T.; Karplus, M. Proteins 1999, 35, 133–152.

(44) Mori, T.; Miyashita, N.; Im, W.; Feig, M.; Sugita, Y. BBA-Biomembranes 2016, 1858, 1635–1651.

(45) Spassov, V. Z.; Yan, L.; Szalma, S. J. Phys. Chem. B 2002, 106, 8726–8738.

(46) Im, W.; Feig, M.; Brooks, C. L., III Biophys. J. 2003, 85, 2900–2918.

(47) Tanizaki, S.; Feig, M. J. Chem. Phys. 2005, 122, 124706.

(48) Panahi, A.; Feig, M. J. Chem. Theory Comput. 2013, 9, 1709–1719.

(49) Dutagaci, B.; Sayadi, M.; Feig, M. J. Comput. Chem. 2017, 38, 1308–1320.

(50) Lazaridis, T. Proteins 2003, 52, 176–192.

(51) Lazaridis, T. Proteins 2005, 58, 518–527.

(52) Zhan, H.; Lazaridis, T. Biophys. J. 2013, 104, 643–654.

(53) Zhan, H.; Lazaridis, T. Biophys. Chem. 2012, 161, 1–7.

(54) Lazaridis, T. J. Chem. Theory Comput. 2005, 1, 716–722.

(55) Nepal, B.; Leveritt, J.; Lazaridis, T. Biophys. J. 2018, 114, 2128–2141.

(56) Iyer, J.; Blankschtein, D. J. Phys. Chem. B 2012, 116, 6443–6454.

(57) Oliver, R. C.; Lipfert, J.; Fox, D. A.; Lo, R. H.; Doniach, S.; Columbus, L. PloS One 2013, 8, e62488.

(58) Faramarzi, S.; Bonnett, B.; Scaggs, C. A.; Hoffmaster, A.; Grodi, D.; Harvey, E.; Mertz, B. Langmuir 2017, 33, 9934–9943.

(59) Barr, A. H. IEEE Computer graphics and Applications 1981, 1, 11–23.

(60) Jaklic, A.; Leonardis, A.; Solina, F., Segmentation and recovery of superquadrics. Springer Science & Business Media: 2013; Vol. 20.

(61) Breen, D. E.; Mauch, S.; Whitaker, R. T. Proc. 1998 IEEE Symp. Volume Visualization 1998, 7–14.

(62) Rahaman, A.; Lazaridis, T. BBA-Biomembranes 2014, 1838, 98–105.

(63) Best, R. B.; Zhu, X.; Shim, J.; Lopes, P. E. M.; Mittal, J.; Feig, M.; Mackerell, A. D., Jr. J. Chem. Theory Comput. 2012, 8, 3257–3273.

(64) Jung, J.; Mori, T.; Kobayashi, C.; Matsunaga, Y.; Yoda, T.; Feig, M.; Sugita, Y. WIREs Comput. Mol. Sci. 2015, 5, 310–323.

(65) Kobayashi, C.; Jung, J.; Matsunaga, Y.; Mori, T.; Ando, T.; Tamura, K.; Kamiya, M.; Sugita, Y. J. Comput. Chem. 2017, 38, 2193–2206.

(66) Brooks, B. R.; Brooks, C. L.; Mackerell, A. D.; Nilsson, L.; Petrella, R. J.; Roux, B.; Won, Y.; Archontis, G.; Bartels, C.; Boresch, S.; Caflisch, A.; Caves, L.; Cui, Q.; Dinner, A. R.; Feig, M.; Fischer, S.; Gao, J.; Hodoscek, M.; Im, W.; Kuczera, K.; Lazaridis, T.; Ma, J.; Ovchinnikov, V.; Paci, E.; Pastor, R. W.; Post, C. B.; Pu, J. Z.; Schaefer, M.; Tidor, B.; Venable, R. M.; Woodcock, H. L.; Wu, X.; Yang, W.; York, D. M.; Karplus, M. J. Comput. Chem. 2009, 30, 1545–1614.

(67) Tieleman, D. P.; van der Spoel, D.; Berendsen, H. J. C. J. Phys. Chem. B 2000, 104, 6380–6388.

(68) Wang, L.; Fujimoto, K.; Yoshii, N.; Okazaki, S. J. Chem. Phys. 2016, 144, 7.

(69) Lomize, M. A.; Lomize, A. L.; Pogozheva, I. D.; Mosberg, H. I. Bioinformatics 2006, 22, 623–625.

(70) Voss, N. R.; Gerstein, M. Nucleic Acids Res. 2010, 38, W555–W562.

(71) Frey, L.; Lakomek, N. A.; Riek, R.; Bibow, S. Angew. Chem. Int. Edit. 2017, 56, 380–383.

(72) MacKenzie, K. R.; Prestegard, J. H.; Engelman, D. M. Science 1997, 276, 131–133.

(73) Trenker, R.; Call, M. E.; Call, M. J. J. Am. Chem. Soc. 2015, 137, 15676–15679.

(74) Da Silva, E. S.; Mulinge, M.; Bercoff, D. P. Retrovirology 2013, 10, 54.

(75) Kliger, Y.; Shai, Y. Biochemistry 1997, 36, 5157–5169.

(76) Hemminga, M. A.; Vos, W. L.; Nazarov, P. V.; Koehorst, R. B. M.; Wolfs, C. J. A. M.; Spruijt, R. B.; Stopar, D. Eur. Biophys. J. 2010, 39, 541–550.

(77) Koehorst, R. B. M.; Spruijt, R. B.; Vergeldt, F. J.; Hemminga, M. A. Biophys. J. 2004, 87, 1445–1455.

(78) Marassi, F. M.; Opella, S. J. Protein Sci. 2003, 12, 403–411.

(79) Papavoine, C. H. M.; Christiaans, B. E. C.; Folmer, R. H. A.; Konings, R. N. H.; Hilbers, C. W. J. Mol. Biol. 1998, 282, 401–419.

(80) Ferrenberg, A. M.; Swendsen, R. H. Phys. Rev. Lett. 1988, 61, 2635–2638.

(81) Kumar, S.; Bouzida, D.; Swendsen, R. H.; Kollman, P. A.; Rosenberg, J. M. J. Comput. Chem. 1992, 13, 1011–1021.

(82) Haass, C.; Selkoe, D. J. Nat Rev Mol Cell Bio 2007, 8, 101–112.

(83) Sugita, Y.; Okamoto, Y. Chem. Phys. Lett. 1999, 314, 141–151.

(84) Miyashita, N.; Straub, J. E.; Thirumalai, D.; Sugita, Y. J. Am. Chem. Soc. 2009, 131, 3438–3439.

(85) Dominguez, L.; Foster, L.; Meredith, S. C.; Straub, J. E.; Thirumalai, D. J. Am. Chem. Soc. 2014, 136, 9619–9626.

(86) Dominguez, L.; Foster, L.; Straub, J. E.; Thirumalai, D. Proc. Natl. Acad. Sci. U.S.A. 2016, 113, E5281–E5287.

(87) Polyansky, A. A.; Volynsky, P. E.; Efremov, R. G. J. Am. Chem. Soc. 2012, 134, 14390–14400.

(88) Xia, B.; Tsui, V.; Case, D. A.; Dyson, H. J.; Wright, P. E. J. Biomol. NMR 2002, 22, 317–331.

(89) Tian, Y.; Schwieters, C. D.; Opella, S. J.; Marassi, F. M. Biophys. J. 2015, 109, 574–585.

(90) Tama, F.; Miyashita, O.; Brooks, C. L., III J. Mol. Biol. 2004, 337, 985–999.

(91) Tanner, D. E.; Chan, K. Y.; Phillips, J. C.; Schulten, K. J. Chem. Theory Comput. 2011, 7, 3635–3642.

(92) Mori, T.; Kulik, M.; Miyashita, O.; Jung, J.; Tama, F.; Sugita, Y. Structure 2019, 27, 161–174.e163.

(93) Miyashita, O.; Kobayashi, C.; Mori, T.; Sugita, Y.; Tama, F. J. Comput. Chem. 2017, 38, 1447–1461.

