## Supplementary for "Implicit micelle model for membrane proteins using super-ellipsoid approximation"

### Minimum distance between a point and super-ellipsoid

In general, minimum distance between an arbitrary point and super-ellipsoid is calculated numerically with iterative minimization schemes.[1] By using polar coordinates $\theta$ and $\phi$, the Cartesian coordinates of a point on the super-ellipsoid can be written as:

$$\begin{aligned} x_s &= a\sin^{m1}\theta\cos^{m2}\phi, \\ y_s &= b\sin^{m1}\theta\sin^{m2}\phi, \\ z_s &= c\cos^{m1}\theta, \end{aligned} \tag{S1}$$

where exponentiation with $m_1$ and $m_2$ is a signed power function (e.g., $\cos^{m1}\theta = \mathrm{sgn}(\cos\theta)|\cos\theta|^{m1}$). The squared distance between the $i$-th solute atom and super-ellipsoid is calculated by

$$\begin{aligned} d_i^2 &= (x_i - x_s)^2 + (y_i - y_s)^2 + (z_i - z_s)^2 \\ &= (x_i - a\sin^{m1}\theta\cos^{m2}\phi)^2 + (y_i - b\sin^{m1}\theta\sin^{m2}\phi)^2 + (z_i - c\cos^{m1}\theta)^2 \, . \\ &= g(\phi,\theta) \end{aligned} \tag{S2}$$

Here, $d_i$ can be considered as a function of $\phi$ and $\theta$. Accordingly, $\phi$ and $\theta$ that give the minimum distance can be numerically calculated, for example, with the simplex algorithm (Nelder-Mead method),[2] which can realize stable minimization and does not require gradient of the target function.

### Derivatives of the solvation free energy

The derivative of the solvation free energy with respect to the atomic position is given by

$$\begin{aligned} \frac{\partial \Delta G_i^{\mathrm{solv}}}{\partial x_i} &= \frac{s}{2\cosh^2(sd_i)}\left(\Delta G_i^{\mathrm{ref,\,water}} - \Delta G_i^{\mathrm{ref,\,cyclohexane}}\right)\frac{x_i - x_s}{d_i} \\ &+ \sum_{j\neq i}\frac{V_j}{2\pi\sqrt{\pi}\lambda_i r_{ij}^2}\exp(-X_{ij}^2)\frac{s}{2\cosh^2(sd_i)}\left(\Delta G_i^{\mathrm{free,\,water}} - \Delta G_i^{\mathrm{free,\,cyclohexane}}\right)\frac{x_i - x_s}{d_i} \\ &+ \sum_{j\neq i}\left\{\frac{\Delta G_i^{\mathrm{free}}}{\pi\sqrt{\pi}\lambda_i r_{ij}^3}\exp(-X_{ij}^2)\left(\frac{1}{r_{ij}} + \frac{X_{ij}}{\lambda_i}\right)V_j + \frac{\Delta G_j^{\mathrm{free}}}{\pi\sqrt{\pi}\lambda_j r_{ij}^3}\exp(-X_{ji}^2)\left(\frac{1}{r_{ij}} + \frac{X_{ji}}{\lambda_j}\right)V_i\right\}(x_i - x_j) \end{aligned} \tag{S3}$$

where $X_{ij} = \{(r_{ij} - R_i)/\lambda_i\}^2$ and $\Delta G^{\mathrm{free}} = f(d)\Delta G^{\mathrm{free,water}} + (1 - f(d))\Delta G^{\mathrm{free,cyclohexane}}$. Since $(x_i - x_s)/d_i$ has a singularity at $d_i = 0$, we replace it with the normal vector $\mathbf{n}$ on the super-ellipsoid surface at $(x_s, y_s, z_s)$:

$$\frac{x_i - x_s}{d_i} = \mathbf{n}_x(x_s, y_s, z_s)\,. \tag{S4}$$

The normal vector $\mathbf{n}$ is calculated by

$$\mathbf{n}(x_s, y_s, z_s) = \frac{\nabla F(x,y,z)}{\| \nabla F(x,y,z) \|} \Big|_{(x_s, y_s, z_s)} . \tag{S5}$$

$(x_s, y_s, z_s)$ is obtained from eq S1, after eq S2 is minimized.

### All-atom MD simulations of pure micelles in explicit solvent

We carried out all-atom MD simulations of DPC, Fos10, Fos14, DHPC, SDS, and LDAO micelles in explicit solvent with various aggregation numbers of surfactant to obtain the IMIC model parameters for pure micelles (Columns 1−3 in Table SI). Typical or previously reported aggregation numbers of the surfactant are 60−80 for DPC,[3] 45−53 for Fos10,[3] 108 for Fos14,[4] 60 for SDS,[5] 69 for LDAO,[6] and 27 or 35 for DHPC.[7, 8] We modeled the systems that contain the surfactant, water, and counter ions by using CHARMM-GUI *micelle builder*,[9] and performed a 100-ns MD simulation (1 step = 2 fs) in the *NPT* ensemble at 300 K and 1 atm for each system using GENESIS.[10, 11] The CHARMM36 force fields were applied to the surfactant molecules.[12] We employed the Langevin thermostat and barostat for temperature and pressure control,[13] and the SHAKE and SETTLE algorithms for bond constraint.[14, 15] For computation of non-bonded interactions, we used the linear $1/R^2$ lookup table method with a cut-off distance of 12 Å,[16] and the particle mesh Ewald method with grid size ~1.2 Å.[17] The trajectories were outputted every 50 ps.

To analyze the three-dimensional mass density profile, we performed a principal component analysis (PCA) for the surfactant hydrocarbon groups in each snapshot to reorient the micelle structure onto the PC1−3 axes, and the atomic coordinates were further projected onto the first octant. Then, we calculated the density profile with a grid size of 0.25×0.25×0.25 $Å^3$. We carried out a least-squares fitting of the function $h\{1 - f(d)\}$ to the profile at the micelle-water interface by randomly changing $a$, $b$, $c$, $m_1$, $m_2$, and $s$ with the Monte Carlo minimization scheme. Here, we used $h$ = 0.745 for DPC, SDS, and LDAO, 0.726 for Fos10, 0.759 for Fos14, and 0.655 for DHPC, which corresponds to the density of *n*-Dodecane, *n*-Decane, *n*-Tetradecane, and *n*-Hexane at 298.15K, respectively.[18, 19]

In Table SI Columns 4−10, we listed the obtained structural parameters $a$, $b$, $c$, $m_1$, $m_2$, $s$, and $V_{\mathrm{surf}}$ for each system. We found that $V_{\mathrm{surf}}$ was not exactly constant, presumably because in larger micelles there are many voids in the micelle center,[20] making the volume of micelle larger. Therefore, to decide the general parameter of each surfactant, we took the averaged value of $V_{\mathrm{surf}}$ among the typical aggregation numbers. From eq 10, we obtained $V_{\mathrm{surf}}$ = 322 $Å^3$ for DPC, SDS, and LDAO, 268 $Å^3$ for Fos10, 379 $Å^3$ for Fos14, and 488 $Å^3$ for DHPC.

### All-atom MD simulations of protein-micelle complexes in explicit solvent

In order to find a general rule for determining the IMIC parameters in the presence of membrane proteins, we performed all-atom MD simulations with explicit solvent for twelve selected membrane-proteins (Hemagglutinin fusion peptide, Integrin β3, Glycophorin A, TMEM14A, AChR β2, BcTSPO, semiSWEET, D3 receptor, KcsA, OmpX, OmpA, and TtoA) in micelles with various aggregation numbers of surfactant (Table SII). We modeled the systems by using the CHARMM-GUI *micelle builder*, and performed a 100−200ns MD simulation (1 step = 2 fs) in the *NPT* ensemble at 300 K and 1 atm using GENESIS. The CHARMM C36 force fields were applied to proteins and surfactants. We employed the Langevin thermostat and barostat for temperature and pressure control and the SHAKE and SETTLE algorithms for bond constraint. For computation of non-bonded interactions, we used the linear $1/R^2$ lookup table method with a cut-off distance of 12.0 Å, and the particle mesh Ewald method with grid size ~1.2 Å.

We analyzed the micelle size and shape around the membrane proteins with the same protocol used for the pure micelles. We calculated the three-dimensional mass density profile for the surfactant hydrocarbon groups with grid size 0.5×0.5×0.5 $Å^3$. Figure S2 shows the mass density profiles of the twelve selected systems. We carried out a least-squares fitting of the function $h\{1 - f(d)\}$ to the micelle-water interface of the mass density profile by randomly changing $a'$, $b'$, $m'_1$, $m'_2$, and $s'$ with the Monte Carlo minimization scheme. Here, $c'$ was fixed to half of the membrane thickness in the OPM database to prevent overfitting. Note that $c'$ was changed in the case of Hemagglutinin. We listed the obtained parameters in the best fitting super-ellipsoid in Table SIII, and also estimated parameters in Table SIV. We found that our estimation was in good agreement with the all-atom MD simulations (Figure S3), where the standard error of the estimate [$\sigma = \sqrt{\sum(X_{\mathrm{MD}} - X_{\mathrm{est}})^2/N}$, where $N = 36$] for $a'$, $b'$, and $m'_1$ was 1.5 Å, 1.3 Å, and 0.08, respectively.

#### MD simulations of GpA in the IMM1 model and explicit lipid bilayers

We performed MD simulations of the glycophorin A (GpA) dimer in DMPC, DOPC, and POPC lipid bilayers. In the IMM1 model, the membrane thickness was set to 25.7 Å for DMPC,[21] 26.8 Å for DOPC,[22] and 28.8 Å for POPC,[21] and the steepness parameter $n = 10$ was used in every case.[23] For the all-atom MD simulations, we modeled the GpA-128DMPC, GpA-128DOPC, and GpA-128POPC complexes by using CHARMM-GUI *membrane builder*,[24] and performed a 200-ns MD run for each system (1 step = 2 fs) in the *NPT* ensemble at 300 K and 1 atm. The CHARMM36 force fields were applied to the proteins and lipids. We used the Langevin thermostat and barostat for temperature and pressure control and the SHAKE and SETTLE algorithms for bond constraint. For computation of non-bonded interactions, we used the linear $1/R^2$ lookup table method with a cut-off distance of 12.0 Å, and the particle mesh Ewald method with grid size of ~1.2 Å.

### MD simulations of the LLP-3 peptide

We carried out MD simulations of the LLP-3 domain in the HIV-1 envelope glycoprotein gp41 with the IMIC and IMM1 models, and compared the structures in these two environments. The amino acid sequence used in this study is ALKYWWNLLQYWSQELKNSAVSL. First, we modeled the ideal α-helical conformation with the backbone dihedral angles of ($\phi$, $\psi$) = (−57.8°, −47.0°), and then placed it at the micelle-water and bilayer-water interface, where the hydrophobic residues of the peptide were facing towards the hydrophobic region of the micelle or bilayer. In the IMIC model, we used the 30DHPC parameters ($a$ = 19.8 Å, $b$ = 14.9 Å, $c$ = 11.4 Å, $m_1$ = 0.95, $m_2$ = 1.0, and $s$ = 0.4), and executed 50 individual runs (500 ns each), starting from random positions and orientations, to search for the optimal position of the peptide on the micelle surface. In the IMM1 model, we used a membrane thickness of 23.1 Å and the steepness parameter $n$ = 10, which mimics a DMPC lipid bilayer. Again, we carried out 8 individual simulations (150 ns each), starting from the same initial coordinates with different random velocities. All simulations were carried out at 300 K with the Langevin thermostat, and the equations of motion were integrated with the leapfrog method with time step 2 fs, where the SHAKE algorithm was employed for bond constraint.

In the IMIC model, the peptide slid on the micelle surface, and moved towards a certain position by changing its orientation. In the IMM1 model, the peptide was stably attached to the membrane surface, and oriented almost parallel to the membrane plane. We analyzed the distribution of the position and orientation of LLP-3 on the micelle and bilayer surfaces. On the micelle, the position was defined by the polar coordinates ($R$, $\Theta$, $\Phi$) of the COM of the peptide, and the orientation was defined by the Euler angles ($\phi$, $\theta$, $\psi$) with respect to the reference α-helix structure located at the origin and along the $X$-axis (Figures S5a−c). Here, to consider the geometrical symmetry, if the COM of the peptide was outside the 1st or 2nd octant, the structure was rotated so that the COM was moved to it. On the bilayer, we analyzed the $Z$-coordinate of the COM of the peptide ($Z_{com}$), and also parts of the Euler angles ($\theta$, $\psi$) (Figure S5a). Analyses for the last 50-ns trajectories showed that the peptide tends to stay around ($R$, $\Theta$, $\Phi$, $\phi$, $\theta$, $\psi$) = (14.8 Å, 71.3°, 52.2°, −25.1°, 37.7°, −55.8°) in the IMIC model, and ($Z_{com}$, $\theta$, $\psi$) = (14.8 Å, 4.0°, 16.2°) in the IMM1 model (Figures S5d−k). Figures 7a and 7b in the main text are the representative snapshots around these positions.

In the PCA described in the main text, we analyzed the backbone heavy atoms from Tyr5 to Ala21 in the last 50-ns trajectories of each MD run, where the conformations with $\Phi$ = 45−60° were used in the IMIC model, and all conformations in the IMM1 model. Note that Figure 7c was illustrated by superimposing the initial structure to the averaged backbone structure used in PCA, and thus the starting points of some vectors in the figure are slightly deviated from the actual atomic positions. In the analysis of the helix-bending angle, we calculated the angle between the N-terminal

(Trp6 to Tyr12) and C-terminal (Ser14 to Ala21) helical axis, each of which were computed from summation of the backbone C=O and H−N vectors.

#### Umbrella sampling of the M13 major coat protein gp8

In the IMIC model, we used a SDS micelle with a typical aggregation number of surfactant ($N_{\mathrm{surf}}$ = 60), where $a'$ = 18.7, $b'$ = 16.2, $c'$ = 15.4, $m'_1$ = 1.0, $m'_1$ = 1.0, and $s'$ = 0.35 were specified. In the IMM1 model we used a membrane thickness of 28.8 Å, corresponding to a POPC lipid bilayer. For the initial structure, we used U-shape of the solution NMR structure (Model 8 in PDB entry: 2CPS). It was embedded into the IMIC/IMM1 models with random orientation while constraining the TM helix (Tyr21−Phe42) inside the hydrophobic core region, and also amphipathic residues near the micelle/bilayer surface. The equations of motion were integrated with the leapfrog method with time step 2 fs, where the SHAKE algorithm was employed for bond constraint. The temperature was controlled with the Langevin thermostat. We used the CHARMM C36m force fields.[25] In the IMIC model, a weak restraint was applied on the distance from the center of mass of the TM domain to the $Z$-axis using a flat-bottom restraint potential (1.5 Å for the switching distance and 1.0 kcal/mol·Å$^2$ for the force constant) to keep the TM domain near the micelle center.

#### REMD simulations of the APP dimer

We carried out REMD simulations of APP in the IMIC and IMM1 models. The amino acid sequence used in this study is DVG$_{25}$SNKGA$_{30}$IIGLM$_{35}$VGGVV$_{40}$IATVI$_{45}$VITLV$_{50}$MLKKK$_{55}$. In the IMIC models, we employed DPC micelles with the surfactant aggregation number of $N_{\mathrm{surf}}$ = 60 or 85. We specified $a'$ = 19.2, $b'$ = 16.7, $c'$ = 16.7, $m'_1$ = 1.0, $m'_1$ = 1.0, and $s'$ = 0.45 for 60DPC, and $a'$ = 22.0, $b'$ = 19.2, $c'$ = 16.7, $m'_1$ = 1.0, $m'_1$ = 1.0, and $s'$ = 0.45 for 85DPC. In the IMM1 model, we used a membrane thickness of 28.8 Å, corresponding to a POPC lipid bilayer, and 32.0 Å. We performed a 100-ns REMD simulation using 16 replicas (1.6 μs in total) for each system. The temperatures were distributed exponentially (287.63, 300.00, 312.67, 325.75, 339.27, 353.26, 367.67, 382.58, 398.00, 413.93, 430.40, 447.42, 465.01, 483.19, 501.97, 521.39 K). The equations of motion were integrated with the leapfrog method with time step 2 fs, where the SHAKE algorithm was employed for bond constraint. The replica exchange was attempted every 2000 step. We used the CHARMM C36m force fields. Since the two helices can be completely separated at higher temperatures in the IMIC and IMM1 models, we applied a weak restraint on the distance from the center of mass of each TM helix to the $Z$-axis using a flat-bottom restraint potential, where we used 12.5 Å for the switching distance between flat and harmonic functions, and 1.0 kcal/mol·Å$^2$ for the force constant. In addition, we applied a flat-bottom restraint for the tilt angle of each TM helix in the IMIC model (45° for the switching angle and 1.0 kcal/mol·rad$^2$ for the force constant) to prevent the helix crossing-angle being larger than 90° at higher temperatures, otherwise the N-terminus of one helix and C-terminus of another helix can interact. In the IMIC model, we further applied a weak restraint on the distance from the center of mass of the whole TM domain to the *Z*-axis using a flat-bottom restraint potential (1.5 Å for the switching distance and 1.0 kcal/mol·$Å^2$ for the force constant) to prevent the protein from large lateral shifting.

In the REMD simulation with the IMIC model, the averaged acceptance ratio in the Metropolis criteria was ~0.24. There was enough overlap between all neighboring pairs of the distributions of the potential energy (Figure S6a). All replicas experienced the temperature at 300 K (Figure S6b), and we observed a random walk in the temperature space (Figure S6c), and also in the potential energy space (Figure S6d). Figure S6e shows the time courses of the Cα-Cα distance between Gly37 in chains A and B, and Figure S6f shows the dihedral angle between the centers of Cα atoms of Lys28−Ala30 in chain A, Met35−Gly37 in chain A, Met35−Gly37 in chain B in 85DPC. As shown in Figure S6g, the two helices were associated and dissociated repeatedly, and the right-handed configurations (negative dihedral angles in Figure S6f) were predominantly sampled compared to the left-handed ones (positive values in Figure S6f). Similar results were obtained in the REMD simulations in 60DPC and IMM1 with the membrane thickness of 28.8 and 32.0 Å. In Figure 9 in the main text, we analyzed the potential of mean force (PMF) by using the weighted histogram analysis method (WHAM),[26, 27] where the first 20-ns REMD trajectories were discarded.

### References


(1) Breen, D. E.; Mauch, S.; Whitaker, R. T. *Proc. 1998 IEEE Symp. Volume Visualization* **1998**, 7-14.

(2) Nelder, J. A.; Mead, R. *Comput. J.* **1965,** *7*, 308-313.

(3) Lipfert, J.; Columbus, L.; Chu, V. B.; Lesley, S. A.; Doniach, S. *J. Phys. Chem. B* **2007,** *111*, 12427-12438.

(4) Oliver, R. C.; Lipfert, J.; Fox, D. A.; Lo, R. H.; Doniach, S.; Columbus, L. *PloS One* **2013,** *8*, e62488.

(5) Gehlen, M. H.; De Schryver, F. C. *J. Phys. Chem.* **1993,** *97*, 11242-11248.

(6) Timmins, P. A.; Leonhard, M.; Weltzien, H. U.; Wacker, T.; Welte, W. *FEBS Lett.* **1988,** *238*, 361-368.

(7) Chou, J. J.; Baber, J. L.; Bax, A. *J. Biomol. NMR* **2004,** *29*, 299-308.

(8) Tausk, R. J. M.; Van Esch, J.; Karmiggelt, J.; Voordouw, G.; Overbeek, J. T. G. *Biophys. Chem.* **1974,** *1*, 184-203.

(9) Cheng, X.; Jo, S.; Lee, H. S.; Klauda, J. B.; Im, W. *J. Chem. Inf. Model.* **2013,** *53*, 2171-2180.

(10) Jung, J.; Mori, T.; Kobayashi, C.; Matsunaga, Y.; Yoda, T.; Feig, M.; Sugita, Y. *WIREs Comput. Mol. Sci.* **2015,** *5*, 310-323.

(11) Kobayashi, C.; Jung, J.; Matsunaga, Y.; Mori, T.; Ando, T.; Tamura, K.; Kamiya, M.; Sugita, Y. *J. Comput. Chem.* **2017,** *38*, 2193-2206.

(12) Klauda, J. B.; Venable, R. M.; Freites, J. A.; O'Connor, J. W.; Tobias, D. J.; Mondragon-Ramirez, C.; Vorobyov, I.; MacKerell, A. D., Jr.; Pastor, R. W. *J. Phys. Chem. B* **2010,** *114*, 7830-7843.

(13) Feller, S. E.; Zhang, Y.; Pastor, R. W.; Brooks, B. R. *J. Chem. Phys.* **1995,** *103*, 4613-4621.

(14) Ryckaert, J.-P.; Ciccotti, G.; Berendsen, H. J. C. *J. Comput. Phys.* **1977,** *23*, 327-341.

(15) Miyamoto, S.; Kollman, P. A. *J. Comput. Chem.* **1992,** *13*, 952-962.

(16) Jung, J.; Mori, T.; Sugita, Y. *J. Comput. Chem.* **2013,** *34*, 2412-2420.

(17) Darden, T.; York, D.; Pedersen, L. *J. Chem. Phys.* **1993,** *98*, 10089-10092.

(18) Grolier, J.-P. E.; Benson, G. C. *Can. J. Chem.* **1984,** *62*, 949-953.

(19) Dubey, G. P.; Tripathi, N.; Bhatia, S. C. *Indian J. Pure Appl. Phys.* **2005,** *43*, 175-179.

(20) Yoshii, N.; Okazaki, S. *Chem. Phys. Lett.* **2006,** *425*, 58-61.

(21) Kučerka, N.; Nieh, M.-P.; Katsaras, J. *BBA-Biomembranes* **2011,** *1808*, 2761-2771.

(22) Pan, J.; Tristram-Nagle, S.; Kučerka, N.; Nagle, J. F. *Biophys. J.* **2008,** *94*, 117-124.

(23) Lazaridis, T. *Proteins* **2003,** *52*, 176-192.

(24) Jo, S.; Lim, J. B.; Klauda, J. B.; Im, W. *Biophys. J.* **2009,** *97*, 50-58.

(25) Huang, J.; Rauscher, S.; Nawrocki, G.; Ran, T.; Feig, M.; de Groot, B. L.; Grubmüller, H.; Mackerell, A. D., Jr. *Nat Methods* **2017,** *14*, 71-73.

(26) Ferrenberg, A. M.; Swendsen, R. H. *Phys. Rev. Lett.* **1988,** *61*, 2635-2638.

(27) Kumar, S.; Bouzida, D.; Swendsen, R. H.; Kollman, P. A.; Rosenberg, J. M. *J. Comput. Chem.* **1992,** *13*, 1011-1021.

**Table SI.** Number of surfactant and water molecules in each pure micelle system and the IMIC model parameters derived from the all-atom MD simulations.

| Surfactant | Num. of mol. | | Parameters in the best-fitting super-ellipsoid | | | | | | |
|---|---|---|---|---|---|---|---|---|---|
| | $N_{surf}$ | $N_{water}$ | $a$ (Å) | $b$ (Å) | $c$ (Å) | $m_1$ | $m_2$ | $s$ | $V_{surf}$ (Å$^3$) |
| DPC (Fos12) | 55 | 13,593 | 18.5 | 16.0 | 14.0 | 0.98 | 1.00 | 0.51 | 319.5 |
| | 60 | 14,820 | 19.2 | 16.5 | 14.3 | 0.98 | 1.00 | 0.50 | 320.2 |
| | 70 | 17,119 | 20.6 | 17.4 | 14.9 | 0.98 | 1.00 | 0.48 | 323.5 |
| | 85 | 21,054 | 22.6 | 18.7 | 15.1 | 0.94 | 1.00 | 0.45 | 326.1 |
| | 100 | 24,106 | 25.5 | 19.6 | 14.9 | 0.92 | 0.99 | 0.41 | 328.7 |
| Fos10 | 45 | 9,232 | 16.8 | 14.1 | 12.0 | 0.99 | 1.00 | 0.55 | 266.2 |
| | 55 | 9,989 | 18.3 | 15.1 | 12.7 | 0.99 | 0.99 | 0.50 | 270.0 |
| | 60 | 10,960 | 18.9 | 15.6 | 13.1 | 1.00 | 1.00 | 0.48 | 269.6 |
| Fos14 | 90 | 21,452 | 23.2 | 20.1 | 16.9 | 0.97 | 1.00 | 0.46 | 373.6 |
| | 100 | 23,256 | 24.3 | 20.9 | 17.2 | 0.95 | 0.99 | 0.44 | 378.7 |
| | 110 | 24,662 | 25.6 | 21.7 | 17.2 | 0.94 | 0.99 | 0.42 | 378.8 |
| SDS | 55 | 16,983 | 19.1 | 15.8 | 13.6 | 1.00 | 0.99 | 0.43 | 313.8 |
| | 60 | 18,566 | 19.6 | 16.3 | 14.1 | 0.99 | 0.99 | 0.41 | 317.6 |
| | 70 | 19,247 | 21.3 | 17.2 | 14.5 | 0.99 | 1.00 | 0.38 | 319.8 |
| | 85 | 23,671 | 24.2 | 18.1 | 14.5 | 0.94 | 1.00 | 0.36 | 324.6 |
| LDAO | 60 | 12,077 | 18.9 | 16.8 | 14.3 | 0.99 | 1.00 | 0.37 | 318.9 |
| | 70 | 13,054 | 21.0 | 17.4 | 14.7 | 0.99 | 1.00 | 0.39 | 323.4 |
| | 85 | 13,908 | 23.2 | 18.6 | 14.9 | 0.95 | 0.99 | 0.37 | 327.9 |
| DHPC | 30 | 15,218 | 19.8 | 14.9 | 11.4 | 0.95 | 0.99 | 0.40 | 486.0 |
| | 35 | 15,706 | 22.0 | 15.4 | 11.3 | 0.90 | 0.98 | 0.38 | 490.4 |
| | 40 | 16,188 | 24.8 | 15.2 | 11.1 | 0.83 | 0.93 | 0.37 | 497.7 |

**Table SII.** Detailed system information of the all-atom MD simulations of protein-micelle complexes.

| Protein | PDB ID | Surfactant | $N_{water}$ | $N_{ion}$ | Time (ns) |
|---|---|---|---|---|---|
| Hemagglutinin | 2KXA | 55 DPC | 20,341 | $2K^+$ | 100 |
| | | 55 SDS | 22,689 | $57Na^+$ | 100 |
| Integrin β3 | 2RN0 | 60 DPC | 26,305 | 0 | 100 |
| | | 60 Fos10 | 14,776 | 0 | 100 |
| Glycophorin A | 1AFO | 60 DPC | 24,641 | $4Cl^-$ | 200 |
| | | 85 DPC | 24,226 | $4Cl^-$ | 200 |
| | | 100 DPC | 25,846 | $4Cl^-$ | 200 |
| | | 85 SDS | 26,950 | $90Na^+$, $9Cl^-$ | 100 |
| TMEM14A | 2LOP | 45 DPC | 28,905 | $8Cl^-$ | 100 |
| | | 70 Fos10 | 20,806 | $8Cl^-$ | 100 |
| | | 70 DPC | 29,972 | $8Cl^-$ | 100 |
| | | 70 Fos14 | 41,544 | $8Cl^-$ | 100 |
| AChR β2 | 2LM2 | 85 DPC | 35,410 | $7K^+$ | 100 |
| | | 85 SDS | 38,656 | $92Na^+$ | 100 |
| | | 85 LDAO | 27,509 | $7K^+$ | 100 |
| BcTSPO | 4RYO | 70 DPC | 33,652 | $4Cl^-$ | 100 |
| | | 85 DPC | 35,510 | $4Cl^-$ | 100 |
| | | 100 DPC | 37,563 | $4Cl^-$ | 100 |
| | | 110 DPC | 39,557 | $4Cl^-$ | 100 |
| semiSWEET | 4RNG | 120 DPC | 32,611 | $8Cl^-$ | 100 |
| | | 135 DPC | 33,555 | $8Cl^-$ | 100 |
| D3 receptor | 3PBL | 135 DPC | 48,075 | $7Cl^-$ | 100 |
| | | 150 DPC | 51,061 | $7Cl^-$ | 100 |
| KcsA | 1R3J | 180 DPC | 49,068 | $8Cl^-$ | 120 |
| | | 220 DPC | 51,200 | $8Cl^-$ | 120 |
| OmpX | 1QJ8 | 60 DPC | 31,040 | $2K^+$ | 200 |
| | | 85 DPC | 30,715 | $2K^+$ | 200 |
| | | 85 SDS | 33,670 | $87Na^+$ | 100 |
| | | 100 Fos10 | 22,050 | $2K^+$ | 100 |
| OmpA | 1BXW | 85 Fos10 | 20,682 | $3K^+$ | 100 |
| | | 85 DPC | 31,245 | $3K^+$ | 100 |
| | | 85 Fos14 | 43,251 | $3K^+$ | 100 |
| TtoA | 3DZM | 70 DPC | 28,918 | $11K^+$ | 100 |
| | | 85 DPC | 30,629 | $11K^+$ | 100 |
| | | 100 DPC | 32,317 | $11K^+$ | 100 |
| | | 110 DPC | 34,322 | $11K^+$ | 100 |

**Table SIII.** Parameters of the super-ellipsoid that was fitted to the micelle-water interface in the mass density profile of the surfactant hydrocarbon group obtained from all-atom MD simulations.

| Protein | Surfactant | $a'$ (Å) | $b'$ (Å) | $c'$ (Å) | $m_1'$ | $m_2'$ | $s'$ |
|---|---|---|---|---|---|---|---|
| Hemagglutinin | 55 DPC | 19.2 | 16.4 | 14.1 | 0.96 | 0.97 | 0.47 |
| | 55 SDS | 19.8 | 16.5 | 13.6 | 0.96 | 0.99 | 0.39 |
| Integrin β3 | 60 DPC | 20.6 | 17.5 | 15.8 | 1.00 | 1.00 | 0.45 |
| | 60 Fos10 | 19.4 | 16.5 | 15.8 | 1.00 | 1.00 | 0.45 |
| Glycophorin A | 60 DPC | 20.2 | 18.4 | 15.95 | 0.95 | 1.00 | 0.44 |
| | 85 DPC | 22.7 | 20.3 | 15.95 | 0.85 | 1.00 | 0.41 |
| | 100 DPC | 23.8 | 20.3 | 15.95 | 0.74 | 0.91 | 0.42 |
| | 85 SDS | 22.7 | 20.1 | 15.95 | 0.82 | 1.00 | 0.35 |
| TMEM14A | 45 DPC | 19.5 | 16.7 | 14.4 | 1.00 | 1.00 | 0.30 |
| | 70 Fos10 | 23.6 | 19.5 | 14.4 | 0.96 | 1.00 | 0.41 |
| | 70 DPC | 24.0 | 20.9 | 14.4 | 0.96 | 1.00 | 0.41 |
| | 70 Fos14 | 25.9 | 20.2 | 14.4 | 0.85 | 0.93 | 0.44 |
| AChR β2 | 85 DPC | 25.4 | 22.4 | 14.6 | 0.72 | 0.98 | 0.41 |
| | 85 SDS | 25.9 | 22.4 | 14.6 | 0.77 | 0.96 | 0.36 |
| | 85 LDAO | 25.8 | 22.0 | 14.6 | 0.64 | 0.99 | 0.43 |
| BcTSPO | 70 DPC | 26.0 | 21.0 | 15.7 | 0.88 | 0.91 | 0.40 |
| | 85 DPC | 26.3 | 22.6 | 15.7 | 0.79 | 0.93 | 0.40 |
| | 100 DPC | 27.9 | 23.1 | 15.7 | 0.73 | 0.94 | 0.45 |
| | 110 DPC | 28.2 | 24.7 | 15.7 | 0.70 | 1.00 | 0.45 |
| semiSWEET | 120 DPC | 29.2 | 25.0 | 17.7 | 0.70 | 1.00 | 0.47 |
| | 135 DPC | 29.9 | 25.8 | 17.7 | 0.67 | 1.00 | 0.46 |
| D3 receptor | 135 DPC | 31.1 | 26.9 | 16.5 | 0.63 | 0.97 | 0.38 |
| | 150 DPC | 31.4 | 28.0 | 16.5 | 0.57 | 1.00 | 0.36 |
| KcsA | 180 DPC | 35.6 | 32.4 | 17.4 | 0.70 | 1.00 | 0.24 |
| | 220 DPC | 38.8 | 34.1 | 17.4 | 0.66 | 1.00 | 0.22 |
| OmpX | 60 DPC | 23.4 | 20.3 | 11.8 | 0.65 | 0.98 | 0.51 |
| | 85 DPC | 26.3 | 22.9 | 11.8 | 0.59 | 1.00 | 0.49 |
| | 85 SDS | 26.3 | 22.2 | 11.8 | 0.55 | 1.00 | 0.39 |
| | 100 Fos10 | 26.6 | 22.8 | 11.8 | 0.57 | 1.00 | 0.48 |
| OmpA | 85 Fos10 | 25.6 | 22.0 | 12.7 | 0.62 | 1.00 | 0.45 |
| | 85 DPC | 26.5 | 23.5 | 12.7 | 0.61 | 0.99 | 0.44 |
| | 85 Fos14 | 27.3 | 23.4 | 12.7 | 0.53 | 0.97 | 0.50 |
| TtoA | 70 DPC | 24.3 | 21.4 | 14.25 | 0.70 | 1.00 | 0.50 |
| | 85 DPC | 25.4 | 22.6 | 14.25 | 0.66 | 0.97 | 0.48 |
| | 100 DPC | 26.6 | 24.2 | 14.25 | 0.63 | 1.00 | 0.52 |
| | 110 DPC | 27.4 | 25.0 | 14.25 | 0.62 | 0.99 | 0.53 |

**Table SIV.** The estimated IMIC model parameters in each test system.

| Protein | $V_{TMD}$ (Å$^3$)[*1] | Surfactant | $a'$ (Å) | $b'$ (Å) | $c'$ (Å)[*2] | $m'_1$ | $m'_2$ | $s'$ |
|---|---|---|---|---|---|---|---|---|
| Hemagglutinin | 0 | 55 DPC[*3] | 18.5 | 16.0 | 14.0 | 1.00 | 1.00 | 0.45 |
| | | 55 SDS[*3] | 19.1 | 15.8 | 13.6 | 1.00 | 1.00 | 0.35 |
| Integrin β3 | 4,108 | 60 DPC | 18.8 | 16.4 | 15.8 | 1.00 | 1.00 | 0.45 |
| | | 60 Fos10 | 17.5 | 15.2 | 15.8 | 1.00 | 1.00 | 0.45 |
| Glycophorin A | 7,251 | 60 DPC | 19.9 | 17.3 | 15.95 | 1.00 | 1.00 | 0.45 |
| | | 85 DPC | 22.5 | 19.6 | 15.95 | 0.97 | 1.00 | 0.45 |
| | | 100 DPC | 23.6 | 20.5 | 15.95 | 0.90 | 1.00 | 0.45 |
| | | 85 SDS | 22.5 | 19.6 | 15.95 | 0.97 | 1.00 | 0.35 |
| TMEM14A | 10,434 | 45 DPC | 20.3 | 17.7 | 14.4 | 1.00 | 1.00 | 0.45 |
| | | 70 Fos10 | 21.6 | 18.8 | 14.4 | 0.94 | 1.00 | 0.45 |
| | | 70 DPC | 22.5 | 19.6 | 14.4 | 0.87 | 1.00 | 0.45 |
| | | 70 Fos14 | 23.4 | 20.4 | 14.4 | 0.82 | 1.00 | 0.45 |
| AChR β2 | 14,584 | 85 DPC | 24.6 | 21.4 | 14.6 | 0.79 | 1.00 | 0.45 |
| | | 85 SDS | 24.6 | 21.4 | 14.6 | 0.79 | 1.00 | 0.35 |
| | | 85 LDAO | 27.5 | 23.9 | 14.6 | 0.67 | 1.00 | 0.45 |
| BcTSPO | 17,587 | 70 DPC | 24.1 | 20.9 | 15.7 | 0.91 | 1.00 | 0.45 |
| | | 85 DPC | 25.0 | 21.8 | 15.7 | 0.85 | 1.00 | 0.45 |
| | | 100 DPC | 26.0 | 22.6 | 15.7 | 0.80 | 1.00 | 0.45 |
| | | 110 DPC | 26.6 | 23.1 | 15.7 | 0.77 | 1.00 | 0.45 |
| semiSWEET | 21,695 | 120 DPC | 27.4 | 23.8 | 17.7 | 0.86 | 1.00 | 0.45 |
| | | 135 DPC | 28.1 | 24.5 | 17.7 | 0.82 | 1.00 | 0.45 |
| D3 receptor | 26,394 | 135 DPC | 29.4 | 25.5 | 16.5 | 0.73 | 1.00 | 0.45 |
| | | 150 DPC | 30.1 | 26.2 | 16.5 | 0.70 | 1.00 | 0.45 |
| KcsA | 43,546 | 180 DPC | 33.8 | 29.4 | 17.4 | 0.65 | 1.00 | 0.45 |
| | | 220 DPC | 35.5 | 30.8 | 17.4 | 0.61 | 1.00 | 0.45 |
| OmpX | 10,801 | 60 DPC | 22.5 | 19.6 | 11.8 | 0.68 | 1.00 | 0.45 |
| | | 85 DPC | 24.8 | 21.6 | 11.8 | 0.59 | 1.00 | 0.45 |
| | | 85 SDS | 24.8 | 21.6 | 11.8 | 0.59 | 1.00 | 0.35 |
| | | 100 Fos10 | 24.6 | 21.4 | 11.8 | 0.60 | 1.00 | 0.45 |
| OmpA | 13,203 | 85 Fos10 | 23.8 | 20.7 | 12.7 | 0.70 | 1.00 | 0.45 |
| | | 85 DPC | 25.0 | 21.7 | 12.7 | 0.65 | 1.00 | 0.45 |
| | | 85 Fos14 | 26.2 | 22.8 | 12.7 | 0.61 | 1.00 | 0.45 |
| TtoA | 14,690 | 70 DPC | 23.7 | 20.6 | 14.25 | 0.82 | 1.00 | 0.45 |
| | | 85 DPC | 24.8 | 21.5 | 14.25 | 0.76 | 1.00 | 0.45 |
| | | 100 DPC | 25.8 | 22.4 | 14.25 | 0.71 | 1.00 | 0.45 |
| | | 110 DPC | 26.5 | 23.0 | 14.25 | 0.69 | 1.00 | 0.45 |

[*1] Calculated from heavy atoms inside the OPM membrane by using the 3V program (Voss and Gerstein, 2010), where the grid spacing of 0.5 Å and probe radius of 3 Å were used.

[*2] Obtained from a membrane thickness in the OPM database (Lomize et al., 2006)

[*3] $a'$, $b'$, and $c'$ were decided based on the pure micelle data (Table SI), since Hemagglutinin is a membrane bound peptide.

**Table SV.** Effective energy and its components (van der Waals, electrostatic, and solvation free energy) of glycophorin A in the IMIC and IMM1 models

| Model | System | $W^d$ | $E_{\mathrm{vdW}}$ | $E_{\mathrm{elec}}$ | $\Delta G_{\mathrm{solv}}$ |
|---|---|---|---|---|---|
| IMIC | 60DPC | 207.9 | −147.6 | −246.2 | −544.3 |
| | 85DPC | 209.1 | −146.9 | −247.0 | −542.7 |
| | 100DPC | 209.8 | −150.1 | −250.9 | −537.8 |
| IMM1 | DMPC | 213.8 | −163.9 | −230.4 | −543.7 |
| | DOPC | 216.1 | −159.1 | −229.8 | −543.1 |
| | POPC | 228.1 | −161.1 | −222.8 | −540.7 |

[d] $W = E_{\mathrm{bond}} + E_{\mathrm{angle,\ UB}} + E_{\mathrm{dihed,\ impr,\ CMAP}} + E_{\mathrm{vdW}} + E_{\mathrm{elec}} + \Delta G_{\mathrm{solv}}$

**Table SVI.** Averaged effective energy and helix crossing-angle of the APP dimer in Basins I-III at 300 K and 447 K.

| Temperature | System | Basin | Effective Energy (kcal/mol) | Helix crossing angle (deg) |
|---|---|---|---|---|
| 300 K | 60 DPC | I | 233.1 | 42.9 |
| | | II | 229.6 | 28.6 |
| | 85 DPC | I | 228.8 | 44.0 |
| | | II | 226.7 | 28.4 |
| | POPC 28.8 Å | I | 245.0 | 44.1 |
| | | II | 247.1 | 28.5 |
| | POPC 32.0 Å | I | 237.6 | 40.3 |
| | | II | 240.8 | 27.1 |
| 447 K | 60 DPC | III | 774.1 | 57.2 |
| | 85 DPC | III | 769.9 | 55.6 |
| | POPC 28.8 Å | III | 786.3 | 49.4 |

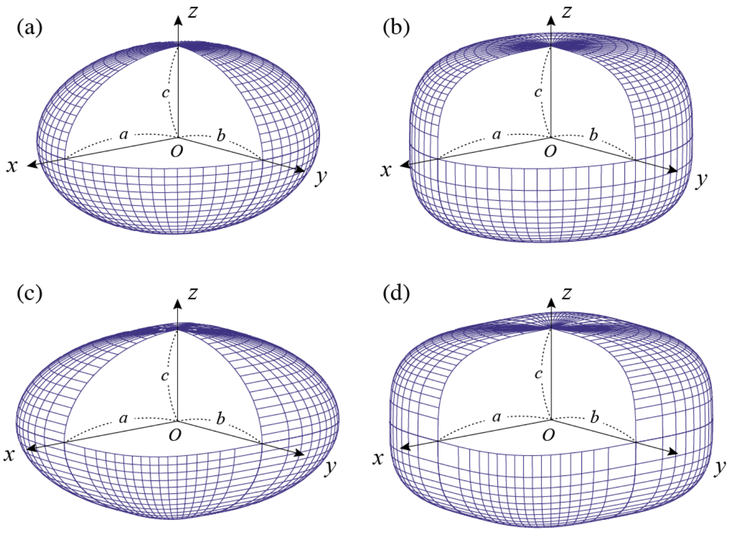


**Figure S1.** Super-ellipsoids represented with the semi-axes $a$, $b$, and $c$ and various $m_1$ and $m_2$. (a) $m_1 = m_2 = 1.0$, (b) $m_1 = 0.65$ and $m_2 = 1.0$, (c) $m_1 = 1.0$ and $m_2 = 0.65$, and (d) $m_1 = m_2 = 0.65$.

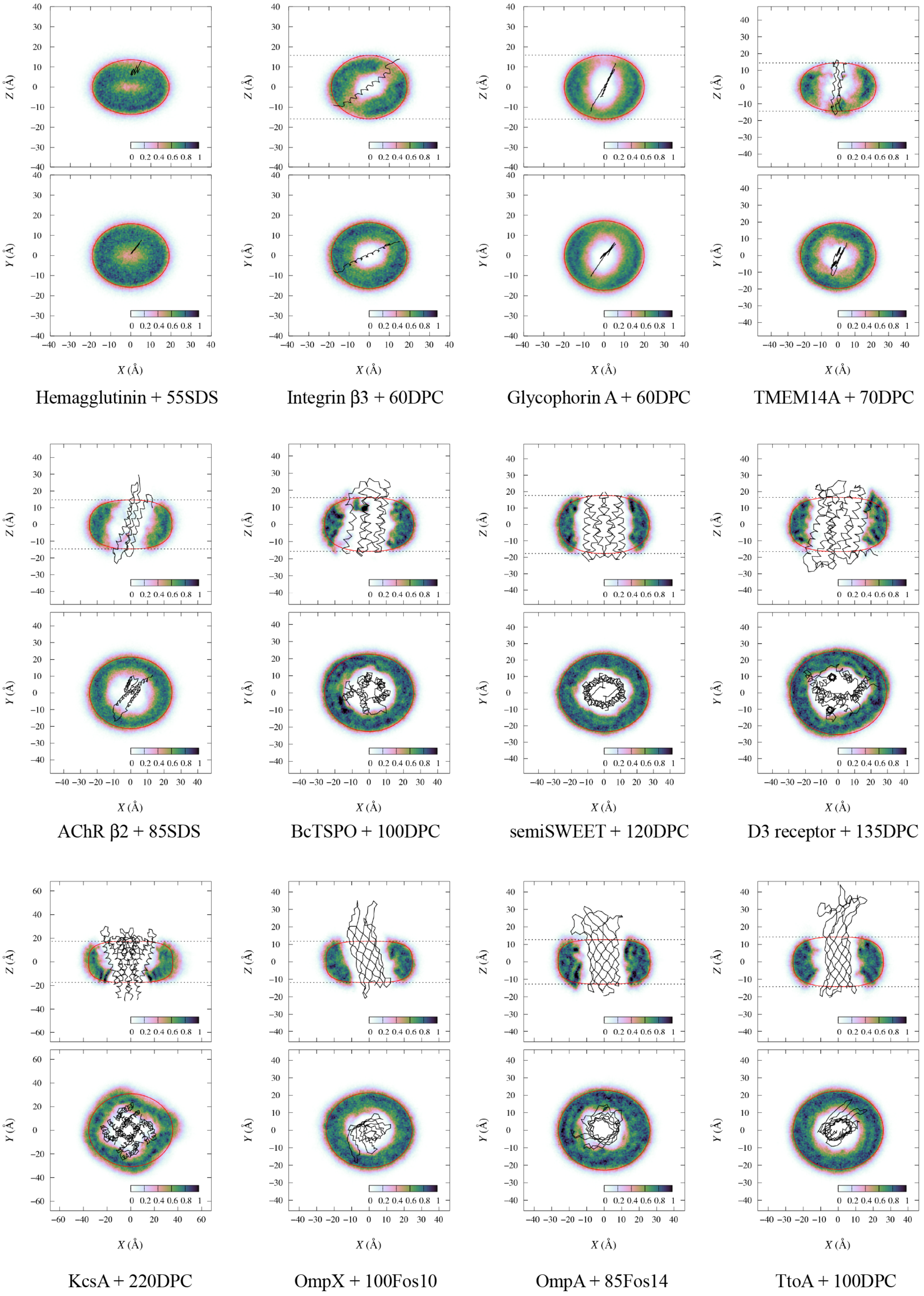


**Figure S2.** Mass density profile in the twelve selected protein-micelle complexes. Black solid line is the protein backbone structure averaged over all snapshots, dotted lines indicate membrane thickness in OPM, and the red line is the micelle-water interface estimated from eqs 7−14. Note that the collapsed protein structure is mainly due to rotation of the protein in the reoriented micelle.

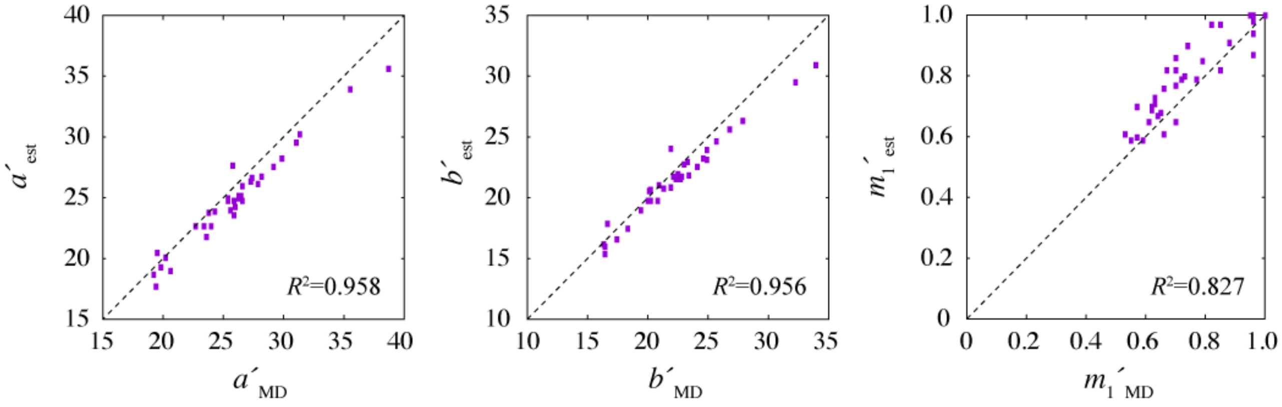


**Figure S3.** Comparison between the parameters $a'$, $b'$, and $m_1'$ estimated from eqs 7−14 (vertical axis; Table SIV) and those obtained in the best fitting super-ellipsoid (horizontal axis; Table SIII).

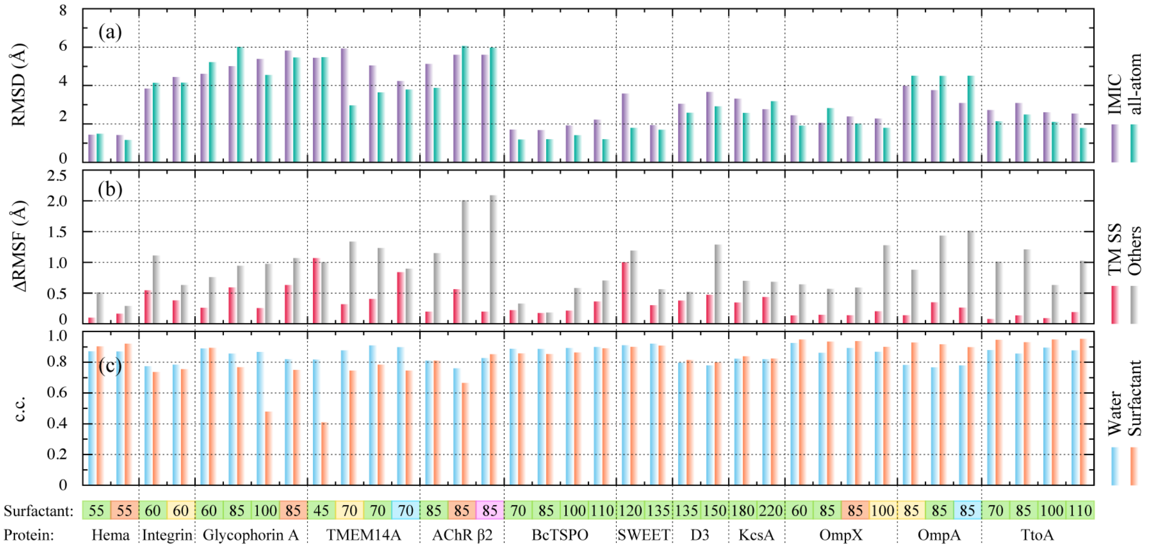


**Figure S4.** Structural and dynamic properties of membrane proteins in the IMIC and all-atom models. Detailed system information is shown at the bottom, where the last line provides the protein name, and the next line above gives the aggregation number of surfactant (yellow: Fos10, green: DPC, blue: Fos14, orange: SDS, and magenta: LDAO). (a) Averaged RMSD for all Cα atoms with respect to the initial structure in the last 50 ns of IMIC (purple) and all-atom (green) simulations. (b) $\Delta$RMSF for the TM secondary structure regions (red) and the other regions (gray). Note that the TM secondary structure region was determined from the OPM database. (c) Correlation coefficient between the SA profiles of IMIC and all-atom models for water-SA (blue) and surfactant-SA (orange).

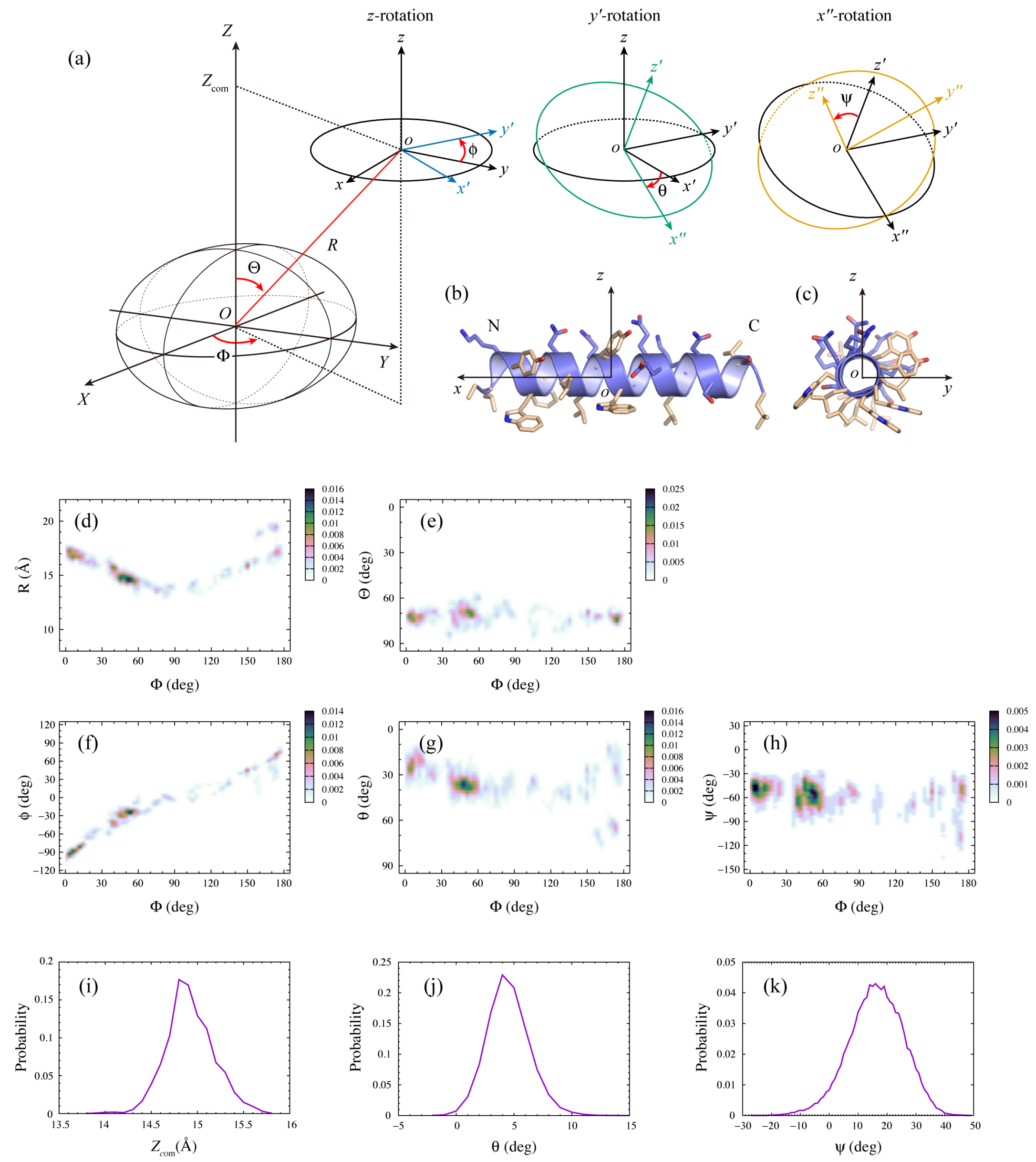

**Figure S5.** Distribution of the position and orientation of the LLP-3 peptide with respect to the IMIC micelle surface and IMM1 bilayer surface. (a−c) Definition of the position and orientation of the peptide using polar coordinates ($R$, $\Theta$, $\Phi$) and the Euler angles ($\phi$, $\theta$, $\psi$). (d−h) Probability distribution of $R$, $\Theta$, $\phi$, $\theta$, and $\psi$ as a function of $\Phi$ in the IMIC model, and (i-k) $Z_{com}$, $\theta$, and $\psi$ in the IMM1 model.

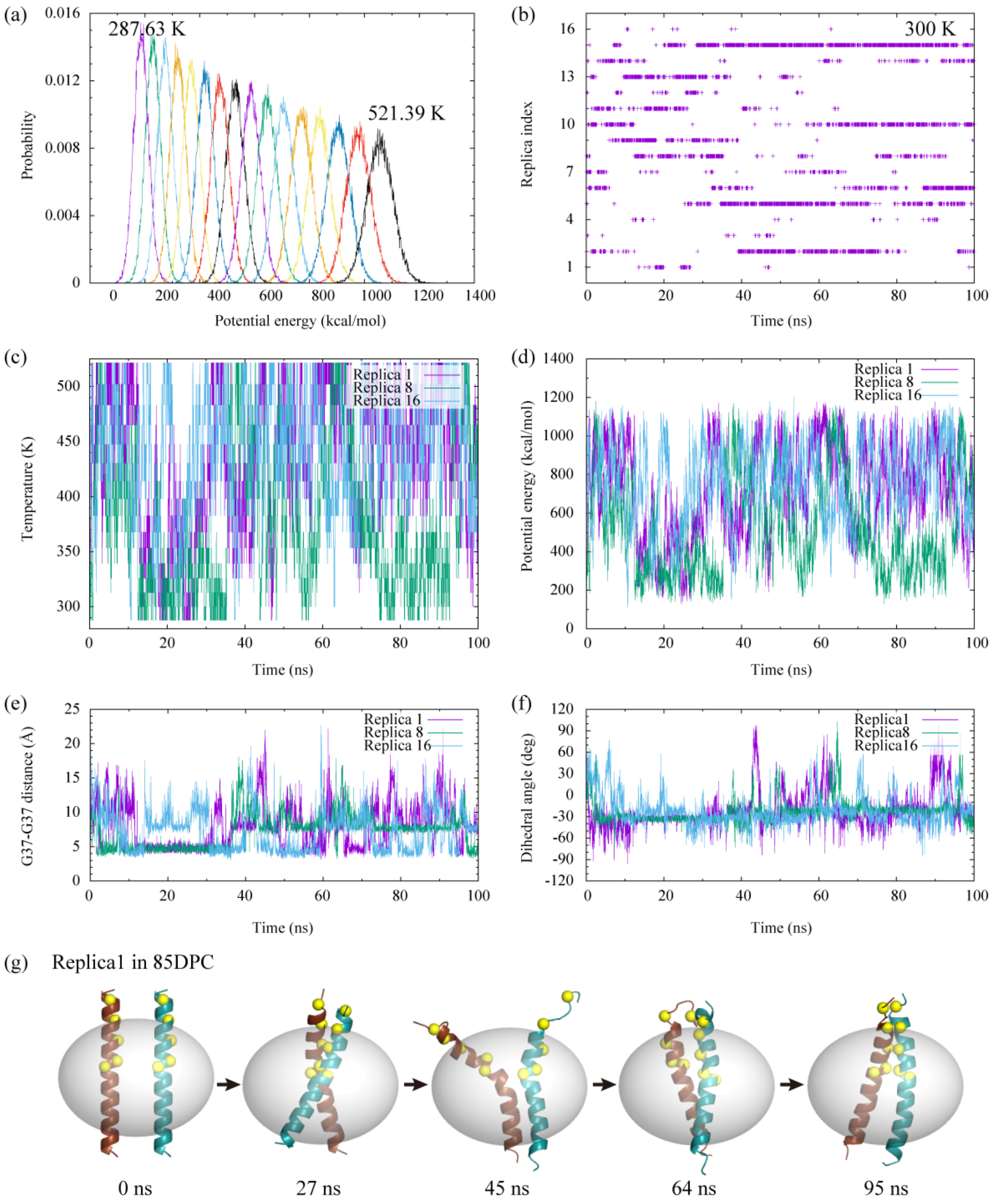


**Figure S6.** REMD simulations of the APP dimer in 85DPC. (a) Probability distribution of the potential energy, (b) time courses of the replica index for 300 K, (c) time courses of the temperature, (d) time courses of the potential energy, (e) time courses of the Gly37-Gly37 Cα distance, and (f) time courses of the dihedral angle between the centers of Cα atoms of Lys28−Ala30 in chain A, Met35−Gly37 in chain A, Met35−Gly37 in chain B, and Lys28−Ala30 in chain B. Positive values indicate left-handed configuration, and negative values are right-handed one. (g) Snapshots of APP in 85DPC in Replica 1.
